# Structural and biochemical basis of FANCI-FANCD2 interdependent ubiquitination

**DOI:** 10.1101/2022.04.07.487446

**Authors:** Kimon Lemonidis, Martin L. Rennie, Connor Arkinson, Viduth K. Chaugule, Mairi Clarke, James Streetley, Helen Walden

## Abstract

The Fanconi Anaemia pathway operates for the repair of interstrand crosslinks and the maintenance of genomic stability upon replication stalling. Di-monoubiquitination of the FANCI-FANCD2 (ID2) complex is a central and crucial step in this pathway. Evidence suggests that FANCD2 ubiquitination precedes FANCI ubiquitination, and that both the FANCD2-ubiquitinated (ID2_Ub_) and the di- monoubiquitinated (I_Ub_D2_Ub_) complex clamp on DNA. However, FANCD2 is deubiquitinated at a faster rate than FANCI, which can result in a FANCI-ubiquitinated ID2 complex (I_Ub_D2). Here, we present a 4.1 Å cryo-EM structure of I_Ub_D2 complex bound to double-stranded DNA. We show that this complex, like ID2_Ub_ and I_Ub_D2_Ub_, is also in the closed ID2 conformation and clamps on DNA. While the target lysine of FANCD2 (K561) is partially buried in the non-ubiquitinated ID2-DNA complex, it becomes fully exposed in the I_Ub_D2-DNA structure, and thus can be ubiquitinated at a faster rate. The I_Ub_D2-DNA complex cannot easily revert to the non-ubiquitinated ID2 state, due to USP1-UAF1-resistance, conferred by the presence of DNA and FANCD2. ID2_Ub_-DNA, on the other hand, can be efficiently deubiquitinated by USP1-UAF1, unless further ubiquitination on FANCI occurs. FANCI ubiquitination also progresses at a faster rate in ID2_Ub_-DNA over ID2-DNA complex, and results in partial DNA-dependent protection from FANCD2 deubiquitination. Taken together, our results suggest that, while FANCD2 ubiquitination promotes FANCI ubiquitination, FANCI ubiquitination in turn maintains FANCD2 ubiquitination by two mechanisms: it prevents excessive FANCD2 deubiquitination within an I_Ub_D2_Ub_-DNA complex, and it enables re-ubiquitination of FANCD2 within a transient, closed-on-DNA, I_Ub_D2 complex.

## Introduction

The Fanconi anaemia (FA) pathway is responsible for repairing interstrand crosslinks (ICLs) and ensuring that genome stability is maintained when replication is stalled (Nalepa & Clapp, 2018). A crucial and central step in this pathway is the mono-ubiquitination of FANCD2 and FANCI on specific lysines (K561 and K523 respectively for human proteins) catalysed by a multi-component ubiquitin ligase (FA-core complex) and the UBE2T ubiquitin-conjugating enzyme (Lemonidis *et al*, 2021). The two ubiquitination events are interdependent, since mutation on either of the two lysines results in greatly impaired in-cell ubiquitination on the other lysine (Smogorzewska *et al*, 2007; Sims *et al*, 2007). Removal of the two ubiquitins, through isopeptide cleavage by the USP1- UAF1 complex, is also required for ICL repair and maintenance of genomic stability (Oestergaard *et al*, 2007; Kim *et al*, 2009; Murai *et al*, 2011).

Several biochemical data (Sato *et al*, 2012; Longerich *et al*, 2014; Rajendra *et al*, 2014; Chaugule *et al*, 2019a; Rennie *et al*, 2020) and recent structural evidence (Wang *et al*, 2021) indicate that FANCD2 is the preferred substrate for ubiquitination and that FANCI ubiquitination likely occurs once FANCD2 has been ubiquitinated. Upon binding to the FA-core-UBE2T, the FANCI-FANCD2 (ID2) complex closes on DNA, and this ID2 remodelling exposes and brings K561 of FANCD2 in proximity to UBE2T’s catalytic cysteine for ubiquitination (Wang *et al*, 2021). The ID2 closure on DNA is maintained upon FANCD2 ubiquitination (Alcón *et al*, 2020; Wang *et al*, 2020; Rennie *et al*, 2020). The resulting ID2_Ub_-DNA complex can be susceptible to USP1-UAF1-mediated deubiquitination. However, further ID2 ubiquitination on FANCI results in enhanced protection of FANCD2’s ubiquitin from USP1-UAF1 action (Rennie *et al*, 2020). Moreover, FANCI appears to be even more resistant to de-ubiquitination than FANCD2, in this DNA-bound di-monoubiquitinated (I_Ub_D2_Ub_-DNA) state (Rennie *et al*, 2020; van Twest *et al*, 2017; Wang *et al*, 2020). Hence, the preferential targeting of FANCD2 for deubiquitination, is likely to result in an ID2 complex that is ubiquitinated on FANCI-only (I_Ub_D2). Currently, we have no information on: i) what conformation such a complex adopts, ii) how does it bind to DNA, iii) how well it supports FANCD2-ubiquitination and iv) how efficiently I_Ub_D2 is protected from deubiquitination.

Providing an answer to such questions would greatly enhance our understanding on how the interdependency in FANCI-FANCD2 *in vivo* ubiquitination (Smogorzewska *et al*, 2007) is encoded at the molecular level, and elucidate the mechanism by which FANCI and FANCD2 ubiquitination (and deubiquitination) are linked. This is clinically relevant too, since FA-pathway modulation is associated with both cancer progression and response to cancer treatment agents. Mutations or overexpression of FA genes and/or USP1, are commonly found in cancers (Niraj *et al*, 2019; Liu *et al*, 2020; García-Santisteban *et al*, 2013; Xu *et al*, 2019). However, and most importantly, FA-gene and/or USP1 deregulation is also frequently associated with chemo-resistance which can be overcome once the expression of the corresponding gene is restored to normal levels (Liu *et al*, 2020; García-Santisteban *et al*, 2013; Xu *et al*, 2019; Lim *et al*, 2018). This suggests that FA- and/or USP1- targeting inhibitors may be beneficial for cancer therapy. USP1, in particular, has been identified as a promising target for cancer-therapy, for a variety of tumours, including: breast (Ma *et al*, 2018; Lim *et al*, 2018; Niu *et al*, 2020; Mussell *et al*, 2020), ovarian (Sonego *et al*, 2019; Lim *et al*, 2018), colorectal (Xu *et al*, 2019), Non-small Cell Lung (Chen *et al*, 2011), bone (Williams *et al*, 2011) and glioma (Ma *et al*, 2019) cancers. Accordingly, there has been growing interest for the development of USP1-UAF1-specific inhibitors (Liang *et al*, 2014; Chen *et al*, 2011). One such USP1- UAF1 inhibitor is currently in Phase I clinical trials, for treatment of advanced solid tumours (KSQ Therapeutics Inc, 2021).

In this work, we show that a transient I_Ub_D2-DNA complex is most likely formed due to significantly faster rate of FANCD2 over FANCI deubiquitination. We further demonstrate that FANCI ubiquitination maintains the closed-on-DNA ID2 conformation when FANCD2 ubiquitination is lost. We lastly show that, in this conformation, FANCD2 ubiquitination is favoured, while FANCI deubiquitination is restricted. Similar to I_Ub_D2-DNA complex having a propensity to transform into an I_Ub_D2_Ub_-DNA complex, the ID2_Ub_-DNA complex also has the propensity to give rise to a di-mono- ubiquitinated complex: this is achieved due to ID2 displaying significantly faster kinetics of FANCI ubiquitination upon FANCD2 ubiquitination. Hence, our results indicate that ubiquitination of either ID2 subunit results in an ID2-DNA clamp that promotes ubiquitination of the other subunit.

## Results

To assess the difference between FANCD2 and FANCI deubiquitination, we assayed I_Ub_D2_Ub_ complex deubiquitination by USP1-UAF1 in a time-course. We found that indeed the rate of FANCI deubiquitination progresses at a much slower rate than FANCD2 deubiquitination (Fig. 1). This suggests that an ID2 complex which is ubiquitinated only on FANCI (I_Ub_D2), may derive from USP1- UAF1-mediated I_Ub_D2_Ub_ deubiquitination. Previous Protein Induced Fluorescence Enhancement (PIFE) assays in our lab showed that fully-ubiquitinated or FANCD2-only-ubiquitinated ID2 complexes display a 10-fold increase in affinity for double-stranded DNA (dsDNA), relative to non-ubiquitinated ID2; however, FANCI-only-ubiquitination results in only a 3-fold enhancement in ID2-DNA affinity (Rennie *et al*, 2020). A possible interpretation for this would be that an I_Ub_D2 complex has a different conformation from ID2 and ID2_Ub_/I_Ub_D2_Ub_, which would allow a different mode of binding to dsDNA. We thus sought to determine the structure of such complex bound to dsDNA, to elucidate how this may differ from ID2_Ub_ and I_Ub_D2_Ub_, and understand how I_Ub_D2 exactly interacts with dsDNA.

**Fig. 1.**
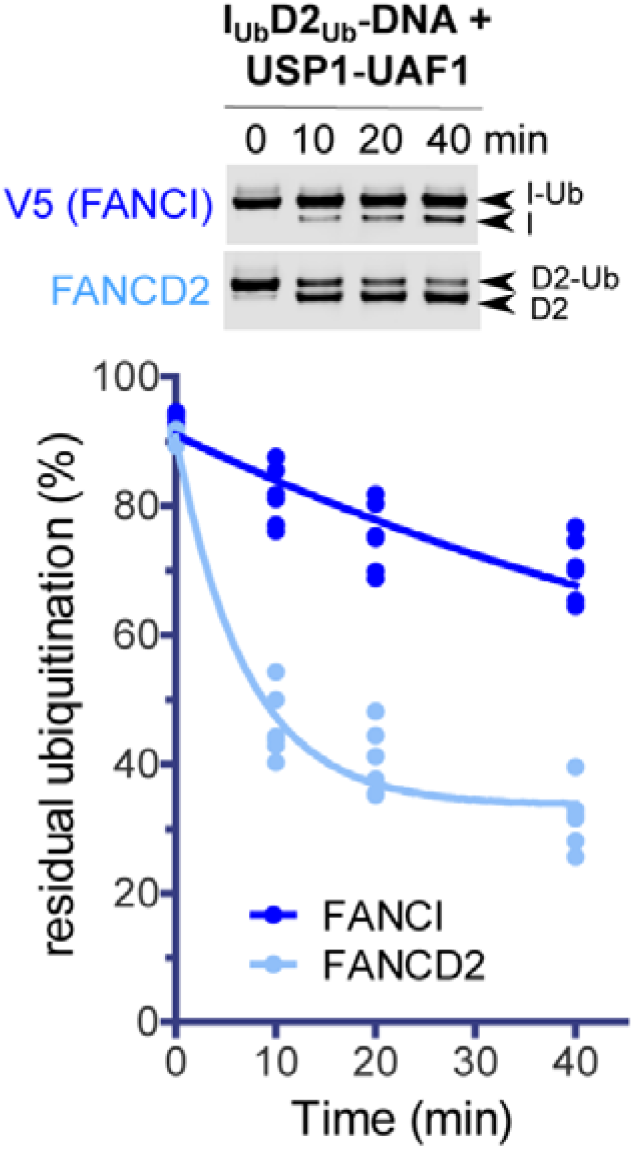
FANCD2 deubiquitination progresses at much faster rate than FANCI deubiquitination. FANCI_Ub_-FANCD2_Ub_-DNA complexes were assembled *in vitro*, and FANCD2_Ub_ and V5-FANCI_Ub_ deubiquitination by USP1- UAF1 (50 nM final) was monitored at room temperature in a time course: at indicative time-points aliquots of each reaction were removed and analysed by western blotting using FANCD2 and V5 antibodies. Experiment was repeated four times and FANCI/FANCD2 ubiquitination levels were calculated following quantification of ubiquitinated and non-ubiquitinated FANCI/FANCD2 bands from the blots. For each protein, all calculated values for all time-points were used for fitting to a one-phase decay model.

To address these questions, we used our *in vitro* reconstitution approach (Arkinson *et al*, 2018; Rennie *et al*, 2020, 2021) to assemble a 1:1:1 I_Ub_D2-DNA complex, from purified I_Ub_, D2 and dsDNA (61 bp long), and subsequently determined its structure by cryo-EM. Such reconstitution approach has been successfully applied to produce ID2_Ub_/I_Ub_D2_Ub_-DNA structures (Rennie *et al*, 2020) and USP1-UAF1-bound ID2_Ub_-DNA structures (Rennie *et al*, 2021). These complexes have been shown to adopt the same closed ID2 conformation as the one observed in ubiquitinated ID2-DNA complexes produced following FA-core-catalysed ID2 ubiquitination (Alcón *et al*, 2020; Wang *et al*, 2020). This suggests that, despite the requirement of FA-core for opening up the ID2 complex for subsequent ubiquitination (Wang *et al*, 2021), ID2 ubiquitination is actually required for both producing and maintaining the final closed ID2 conformation. Hence, we reasoned that our *in vitro* assembled complex would also be structurally indistinguishable from a complex produced through removal of FANCD2’s ubiquitin from I_Ub_D2_Ub_. Our final I_Ub_D2-DNA map, made of 139,601 image particles, was at 4.14Å global resolution and had a local resolution ranging from 2.8Å to 13.9Å (Fig. EV1A-E; Table 1). By 2D classification we also obtained few smaller-sized-particle 2D class averages (4 classes; 62,961 particle images in total), likely corresponding to dissociated monomeric proteins (Fig. EV2). Using the structure of I_Ub_D2_Ub_-DNA (Wang *et al*, 2020)(PDB: 6VAE) as initial model (but with the ubiquitin conjugated to FANCD2 removed) for refinement, we obtained an atomic model of the I_Ub_D2-DNA structure at 4.1Å resolution (Table 1; Fig. 2A and EV1E). Our maps had several well- resolved regions for modelling, like the one surrounding and including FANCI’s K523 isopeptide linkage with G76 of ubiquitin (Fig. EV1F) and the region of FANCI-FANCD2 C-termini interaction (Fig. EV1G). Many FANCI and FANCD2 loops, as well as the FANCI N-terminus (region corresponding to the first 170 aa) had very poor density and were thus unmodelled (Fig 2A). Modelled regions of relatively poor density included the dsDNA (Fig. EV1H), the central region of FANCD2 and an N- terminal part of FANCI (Fig. EV1E). Interestingly, we found that I_Ub_D2 has the same closed-on-DNA conformation as I_Ub_D2_Ub_ and ID2_Ub_ (Fig. 2A). We hypothesized that the apparent lower than expected enhancement of ID2-DNA affinity upon FANCI ubiquitination measured before, may had been due to I_Ub_ dissociating from D2 at low concentrations. Indeed, previous PIFE assays showed that at lower concentrations of I_Ub_D2 (<100 nM) there had been negligible protein-binding induced fluorescence enhancement of labelled DNA, while at higher protein concentrations (>500 nM), the I_Ub_D2 -binding induced fluorescence enhancement of labelled DNA, had been comparable to that achieved with I_Ub_D2_Ub_ and ID2_Ub_ complexes (Rennie *et al*, 2020). Hence, to ensure complex formation at low FANCI concentrations, we performed dsDNA-binding PIFE assays for ID2 and I_Ub_D2 as before (Rennie *et al*, 2020), but this time we titrated only FANCI (ubiquitinated or not), while having a constant high concentration of FANCD2 - equal to the maximum concentration of FANCI used. With such set-up, our PIFE assays revealed a 20-fold increase in ID2 affinity for dsDNA when FANCI was ubiquitinated, whereas FANCD2 on its own had negligible binding to dsDNA (Fig. 2B). The above indicate that FANCI-ubiquitination is responsible for maintaining the clamping of the ID2 complex on DNA, when FANCD2 ubiquitination is lost.

**Table 1.** Cryo-EM data collection and processing, and subsequent model building and refinement.

| Data Collection and Processing |  |
| --- | --- |
| Magnification | 120,000x |
| Voltage (kV) | 300 |
| Electron exposure (e/Å<sup>2</sup>) | 45.2 or 46.8 |
| Pixel size (Å) | 1.023 |
| Defocus range (µm) | 0.5-3.8 |
| Symmetry imposed | C1 |
| Initial number of images (automated picking) | 7,376,277 |
| Final particle images | 139,601 |
| Map resolution (Å) at FSC=0.143 | 4.1 |
| Map resolution range (Å) at FSC=0.143 | 2.8-13.9 |

| Refinement and Validation |  |  |
| --- | --- | --- |
| Initial model used (PDB code) |  | 6VAE |
| Model resolution (Å) | Resolution at FSC=0.143 | 4.1 |
|  | Resolution at FSC=0.5 | 4.4 |
| Model composition | Non-hydrogen atoms | 19,458 |
|  | Protein residues | 2,297 |
|  | DNA residues | 54 |
| All-Atom contacts | Clash score | 3.24 |
| Bonds RMSD | Bond Lengths (Å) | 0.005 |
|  | Bond Angles (°) | 0.81 |
| Ramachandran plot | Favoured (%) | 97.34 |
|  | Allowed (%) | 2.66 |
|  | Outliers (%) | 0 |
|  | Rama-Z score (RMSD) | 0.76 |
| Protein Geometry | MolProbity score | 1.24 |
|  | Rotamer outliers (%) | 0 |
|  | Cβ outliers (%) | 0 |
|  | Twisted peptides (%) | 0 |
|  | CaBLAM outliers (%) | 0.63 |
| B-factors (Å<sup>2</sup>) | Protein | 127.49 |
|  | Nucleotide | 306.24 |
| Map-model correlation coefficients | CC<sub>MASK</sub> | 0.74 |
|  | CC<sub>BOX</sub> | 0.78 |

**Fig. 2.**
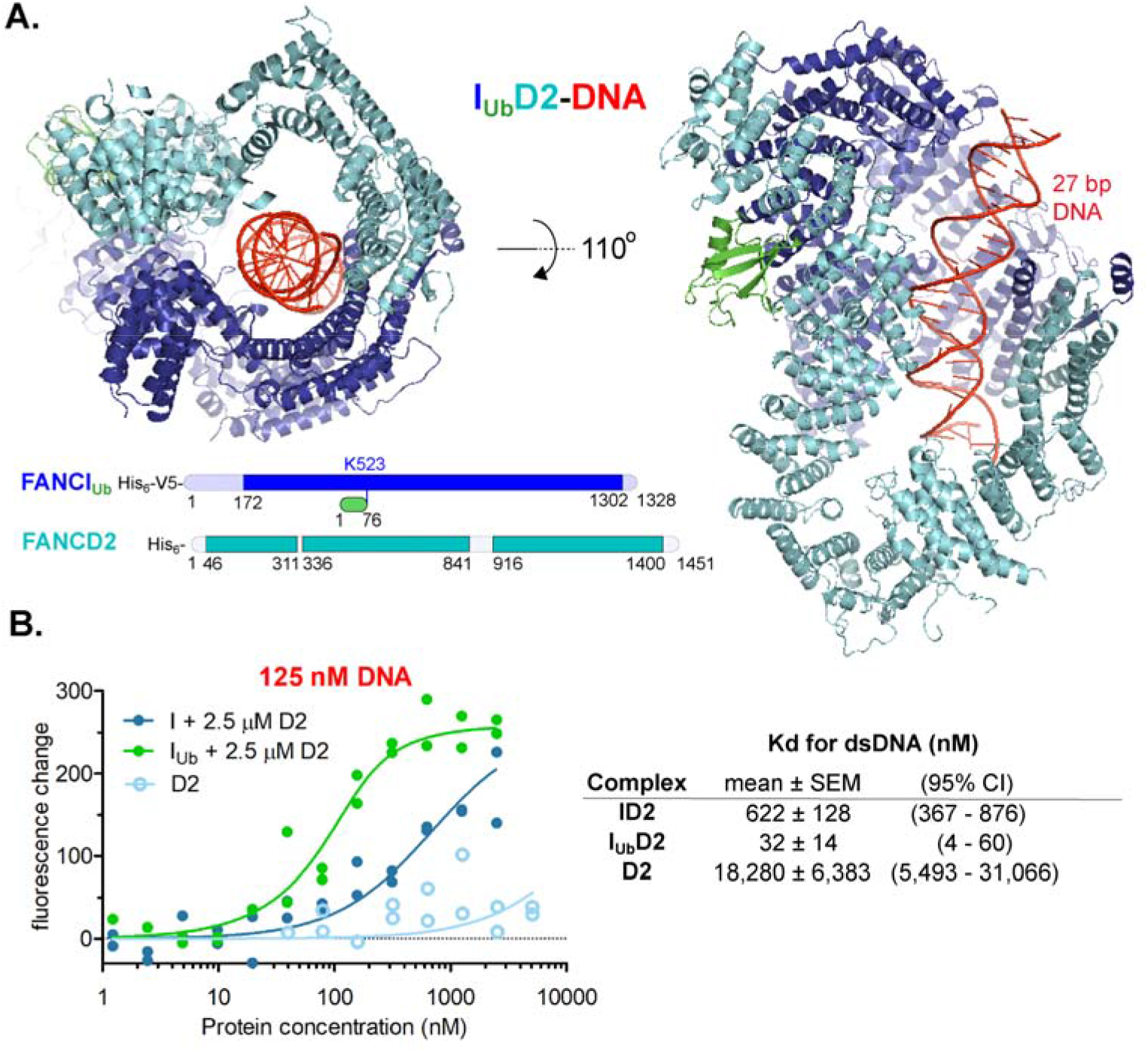
FANCI_Ub_-FANCD2 complex is a DNA clamp. **A**. FANCI_Ub_-FANCD2 (I_Ub_D2) structure bound to double-stranded DNA. The structure was determined by cryo-EM, using a 4.1Å global resolution map. Two different views of the structure are shown. Unmodelled regions due to poor density that extend 20 amino-acid stretches are indicated at the bottom. **B**. FANCI-ubiquitinated ID2 complex displays increased affinity to double-stranded DNA (dsDNA) compared to non-ubiquitinated ID2 complex. *Left*: Fluorescent changes of IRDye700-labeled 32 bp DNA (125 nM) when incubated with FANCD2 (D2; 2.5 µM) and increasing concentrations (ranging from 1.2 nM to 2.5 µM) of FANCI (I) or ubiquitinated FANCI (I_Ub_). As a control, fluorescent changes of IRDye700-labelled DNA (125 nM) when incubated with increasing concentrations of FANCD2 (ranging from 40 nM to 5 µM) were monitored as well. For each protein/complex, the experiment was conducted twice and all data points from the two experiments were used for fitting of a one-site binding model. *Right*: Apparent ID2, I_Ub_D2 and D2 Kd values (and associated uncertainties, all in nM) for dsDNA measured from model fitting.

The overall conformation of the I_Ub_D2 complex is very similar to that of I_Ub_D2_Ub_ (Fig. EV2A). The most noticeable differences are: i) slight movements of FANCD2 and FANCI helices in the region where FANCD2’s ubiquitin interacts with FANCI; and ii) the high level of disorder in the FANCI N-terminus proximal to that region (residues 1-170), upon loss of FANCD2 ubiquitination (Fig. EV2B-C).

While both I_Ub_D2_Ub_-DNA (EMD-21138) and ID2_Ub_-DNA (EMD-21139) maps display relatively poor density for FANCI N-terminus (Wang *et al*, 2020), there is virtually no density for that part in both our locally-filtered and Phenix-auto-sharpened map (Fig EV1E and Fig. 3A, respectively). Lack of density in the N-terminus of FANCI has been also observed before, upon extraction of FANCD2’s ubiquitin by USP1 (Rennie *et al*, 2021). We thus propose that the high level of disorder in the FANCI N-terminus is a direct consequence of the loss of binding between the ubiquitin of FANCD2 and the N-terminus of FANCI. Similarly, reduced density for the N-terminus of FANCD2 has been previously observed in closed state ID2 conformations in which there is no ubiquitin conjugated to FANCI, like in the ID2_Ub_-DNA (EMD-21139) map (Wang *et al*, 2020) (Fig. 3A; *right*) and USP1-UAF1-ID2_Ub_-DNA (EMD-11934) map (Rennie *et al*, 2021).

**Fig. 3.**
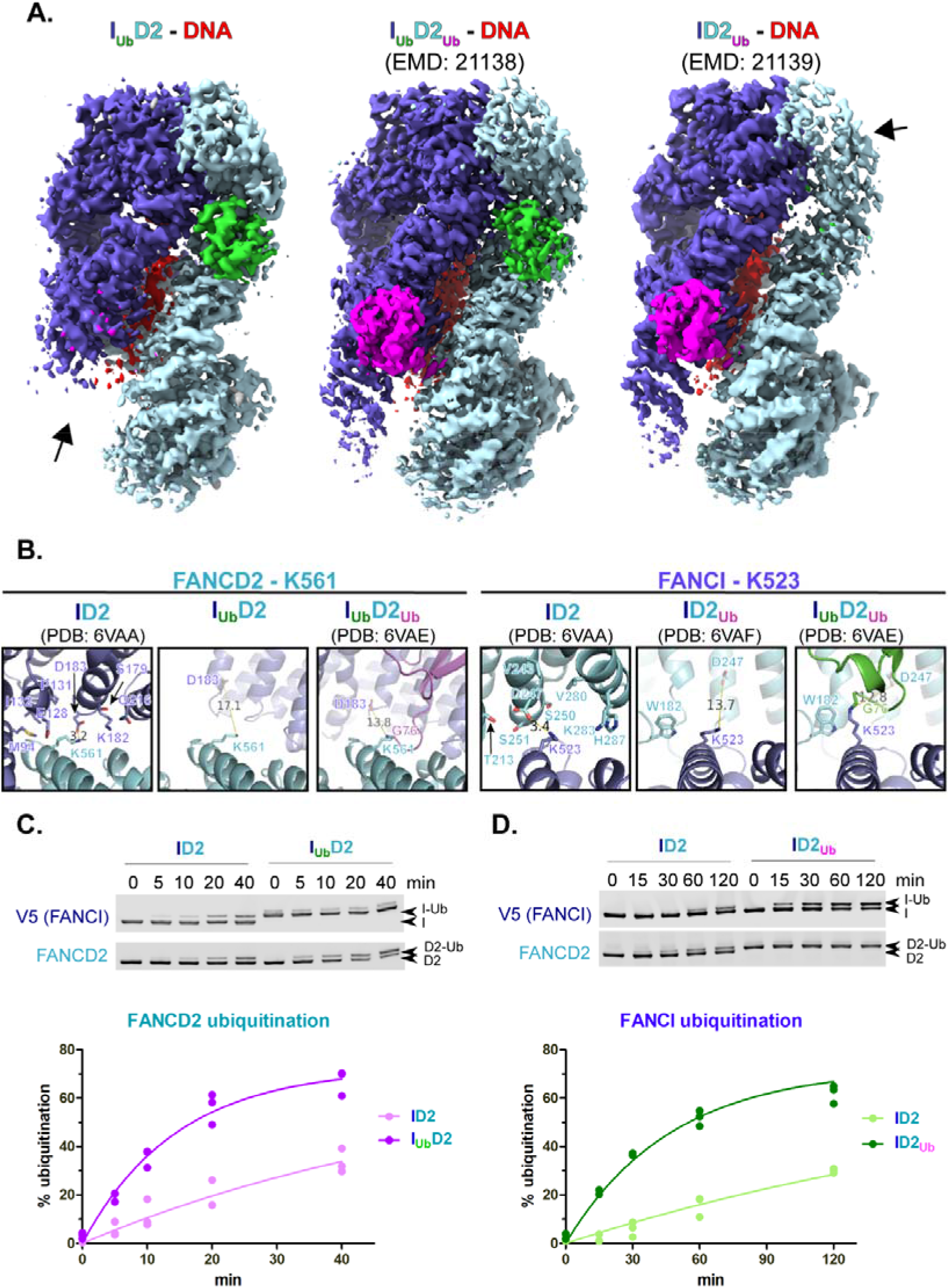
Ubiquitination of either of the two ID2 subunits enhances ubiquitination of the other. **A**. Comparison of cryo-EM density distribution among I_Ub_D2-DNA (Phenix-auto-sharpened map), I_Ub_D2_Ub_-DNA (EMD: 21138) and ID2_Ub_-DNA (EMD: 21138) maps. I_Ub_D2-DNA and ID2_Ub_-DNA maps, as well as I_Ub_D2_Ub_-DNA model (PDB: 6VAE) were aligned to I_Ub_D2_Ub_-DNA in ChimeraX. A different colour was applied for each of the protein chains of I_Ub_D2_Ub_-DNA model (FANCI: slate blue, Ubiquitin-on- FANCI: green, FANCD2: cyan, Ubiquitin-on-FANCD2: magenta), while DNA was coloured red. Then each map was coloured according to nearby (within 6Å) residue colours. Contour levels were adjusted (I_Ub_D2-DNA: 6.21, I_Ub_D2_Ub_-DNA: 0.0194 and ID2_Ub_-DNA: 0.0162) to achieve comparable volumes among all displayed maps (ranging from 8.6 to 9.4 × 10^4^ Å^3^). Arrows indicate regions of poorer density (in I_Ub_D2-DNA and ID2_Ub_-DNA maps) relative to other regions of the map, as well as to equivalent positions in ID2_Ub_-DNA map. **B**. Both K561 of FANCD2 and K523 of FANCI become more accessible upon ubiquitination of the other ID2 subunit. Structural comparison of relative accessibility of FANCD2-K561, upon FANCI ubiquitination (*left panel*), and of FANCI-K523, upon FANCD2 ubiquitination (*right panel*). The relative positions of these lysines upon conjugation with ubiquitin, are also shown for comparison. Residues of the other ID2 subunit within 8 Å distance from the epsilon-amino-group of the corresponding lysine are indicated as sticks. The distance to the nearest residue is shown prior and upon ubiquitination of the other ID2 subunit. In either case this increases, upon ubiquitination of the other subunit, further than 10 Å. **C-D**. ID2 ubiquitination on FANCI results in increased rate of FANCD2 ubiquitination (B), whereas ID2 ubiquitination on FANCD2 results in increased rate of FANCI ubiquitination (C). Protein complexes were assembled *in vitro* on ice in the presence of dsDNA (32 bp) and their *in vitro* ubiquitination at 30° C was subsequently monitored in a time-course: at indicative time-points aliquots of the reaction were removed and analysed by western blotting using FANCD2 and V5 antibodies (*Top*). For each protein complex, data-points from three replicate experiments were used in fitting to a one-phase association model (*Bottom*).

We reasoned that the relative disorder in the N-terminal regions of FANCI or FANCD2, in I_Ub_D2-DNA or ID2_Ub_-DNA complexes respectively, might be crucial for ubiquitination of FANCD2 or FANCI, correspondingly. Despite the slight movements of I_Ub_D2’s FANCI and FANCD2 towards the region where FANCD2-conjugated ubiquitin would be (Fig. EV2B-C), FANCD2’s K561 is fully accessible for ubiquitination, when compared to its position in the ID2-DNA complex (79 Å^2^ buried) (Fig. 3B; Fig. EV3). Similarly, several FANCD2 residues in proximity to FANCI’s K523 (in ID2-DNA complex) are positioned further away upon FANCD2 ubiquitination, and hence FANCI’s K523 becomes more accessible for ubiquitination in the ID2_Ub_-DNA complex than in the ID2-DNA complex (Fig. 3B; Fig. EV3). These observations led us to the hypothesis that ubiquitination of either of the two subunits of ID2 (FANCI or FANCD2) actually favours ubiquitination of the other subunit. Indeed, time-course ubiquitination assays revealed that: FANCD2 ubiquitination is stimulated when FANCI is already ubiquitinated; and similarly, FANCI ubiquitination is stimulated when FANCD2 is already ubiquitinated (Fig. 3C). These results may partially explain the *in vivo* interdependency in FANCI and FANCD2 ubiquitination observed before (Smogorzewska *et al*, 2007; Sims *et al*, 2007).

Previous work has shown that DNA is required for efficient protection of both FANCI and FANCD2 against USP1-UAF1 mediated deubiquitination (van Twest *et al*, 2017; Arkinson *et al*, 2018). Focusing on FANCD2 deubiquitination, we have previously found that FANCI ubiquitination (but not FANCI itself) is additionally required for restricting FANCD2 deubiquitination, but has no effect in protecting from FANCD2 deubiquitination, in the absence of dsDNA (Arkinson *et al*, 2018). We wondered whether a similar mechanism exists for protection of FANCI ubiquitination: i.e. the very slow FANCI deubiquitination in the I_Ub_D2_Ub_-DNA complex (Fig. 1) may be due I_Ub_ being protected from USP1-UAF1-mediated deubiquitination, when associating with both D2_Ub_ and DNA. Alternatively, the presence of simply dsDNA, or FANCD2 (irrespective of ubiquitination status) and dsDNA, may hinder USP1-UAF1 from targeting FANCI’s ubiquitin. To test those possibilities we performed USP1-UAF1 deubiquitination assays with either isolated FANCI/FANCD2 proteins (I_Ub_ or D2_Ub_), or differentially ubiquitinated ID2 complexes (I_Ub_D2, I_Ub_D2_Ub_ or ID2_Ub_), in the presence or absence of dsDNA (Fig. 4A). We observed that dsDNA significantly protects against FANCI deubiquitination, whether I_Ub_ is in isolation, or in complex with D2/D2_Ub_. Moreover, the protective role of DNA against FANCI deubiquitination, is further enhanced when I_Ub_ is in complex with FANCD2, and this enhancement was irrespective of FANCD2 ubiquitination status (Fig. 4A-B). This suggests that both DNA and FANCD2 are required for maximal protection of I_Ub_ against USP1-UAF1 deubiquitination. In agreement with what we observed before (Arkinson *et al*, 2018; Rennie *et al*, 2020), the presence of FANCI did not affect FANCD2 deubiquitination, which was nearly complete in our reaction conditions, whether FANCI was present or not (Fig. 4A). However, the inclusion of ubiquitinated FANCI in our reactions, restricted to some extent FANCD2 deubiquitination, when DNA was also present (Fig. 4A-B).

**Fig. 4.**
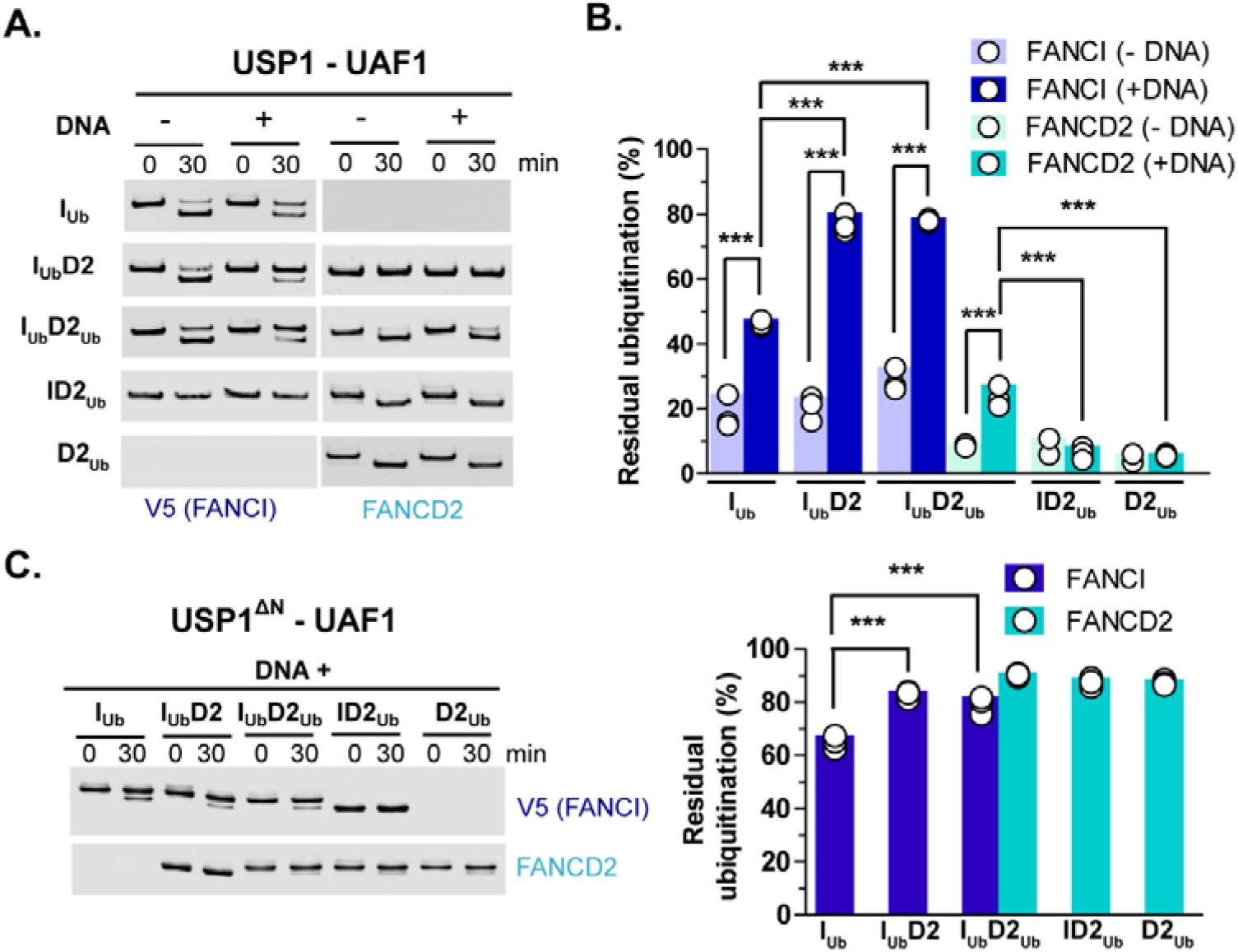
DNA and FANCD2 protect against FANCI deubiquitination. **A**. USP1-UAF1-mediated deubiquitination of V5-FANCI and FANCD2 was assessed in the absence or presence of DNA (51 bp), when ubiquitinated versions of these proteins were in isolation (I_Ub_ and D2_Ub_) or within singly/doubly ubiquitinated ID2 complexes (I_Ub_D2, I_Ub_D2_Ub_ and ID2_Ub_). At indicated time-points aliquots of each reaction were removed and analysed by western blotting using FANCD2 and V5 antibodies. **B**. Residual FANCI and FANCD2 ubiquitination following USP1-UAF1 treatment for 30 min at room temperature. Experiments shown in A were performed in triplicate, apart from ID2_Ub_ and D2_Ub_ deubiquitination in the absence of DNA, which were performed twice (and were thus excluded from statistical analysis). Replicate residual ubiquitination values and statistically significant changes (one-way ANOVA test with Bonferroni correction) are shown. ***p<0.001. **C**. Deletion of N-terminus (ΔN) of USP1 (residues 1-54) results in greatly reduced FANCD2 deubiquitination. Assays were performed in triplicate as in A, but all reactions contained DNA. *Left*: Western blotting of reaction products at zero and 30 minutes using FANCD2 and V5 antibodies. *Right*: Residual FANCI and FANCD2 ubiquitination following USP1-UAF1 treatment for 30 min. Replicate residual ubiquitination values and statistically significant changes (one-way ANOVA test with Bonferroni correction) are shown. ***p<0.001.

Relative to FANCI, FANCD2 is efficiently deubiquitinated. This is achieved thanks to a USP1 N-terminal region (proximal to its USP domain) specifically targeting FANCD2 (Arkinson *et al*, 2018). Indeed, when deubiquitination occurred under same conditions, but with a USP1 having this N-terminal region (first 54 amino-acids) deleted (USP1ΔN), FANCD2 deubiquitination was nearly abolished, whether D2_Ub_ was in isolation or in complex with I/I_Ub_. The USP1 substitution with the USP1ΔN mutant in our assays, however, did not greatly affect FANCI deubiquitination (Fig. 4C). Moreover, the presence of either D2 or D2_Ub_, provided I_Ub_-deubiquitination protection from USP1ΔN-UAF1 (Fig. 4C), similarly to what observed with wild-type USP1-UAF1 complex (Fig. 4A-C). The above suggest that the interaction of FANCD2’s ubiquitin with FANCI (in the I_Ub_D2_Ub_-DNA complex) does not protect against FANCD2 deubiquitination, as the latter can be efficiently achieved by a mechanism that involves USP1’s N-terminus binding to FANCD2. Crucial for such binding are residues R22 and L23 of USP1, as predicted by AlphaFold modelling of FANCD2 interaction with USP1’s N-terminus (Rennie *et al*, 2022), and further supported by deubiquitination assays of respective USP1 alanine mutants towards FANCD2, FANCI and PCNA (Arkinson *et al*, 2018).

Whereas the FANCI interaction with the ubiquitin conjugated to FANCD2 has no protective role against USP1-UAF1 mediated FANCD2 deubiquitination, the interaction of FANCI’s ubiquitin with FANCD2 (in I_Ub_D2-DNA and I_Ub_D2_Ub_-DNA complexes) efficiently protects against FANCI deubiquitination. In each case, ubiquitin’s hydrophobic I44 patch is involved in interaction with the other ID2 subunit; however, the ubiquitin conjugated to FANCI forms a more extended interface with FANCD2 (Wang *et al*, 2020)(Fig. 5A). This extended interface is formed predominantly via hydrophobic interactions of residues H209 (α10 helix), V243, P244 and D247 (α13 helix) of FANCD2 with residues T9 and K11 of ubiquitin, and is further stabilized by hydrogen bonding between ubiquitin’s T9 and R74 with FANCD2’s H209 and S251 (or E217), respectively (Fig. 5A). The residues of FANCD2 predicted to be involved in this extended interface show a high level of conservation among vertebrate species (Fig. 5B). Hence, the above interactions may be crucial for the maintenance of ID2 ubiquitination (and therefore of ID2 clamping on DNA) in vertebrates.

**Fig. 5.**
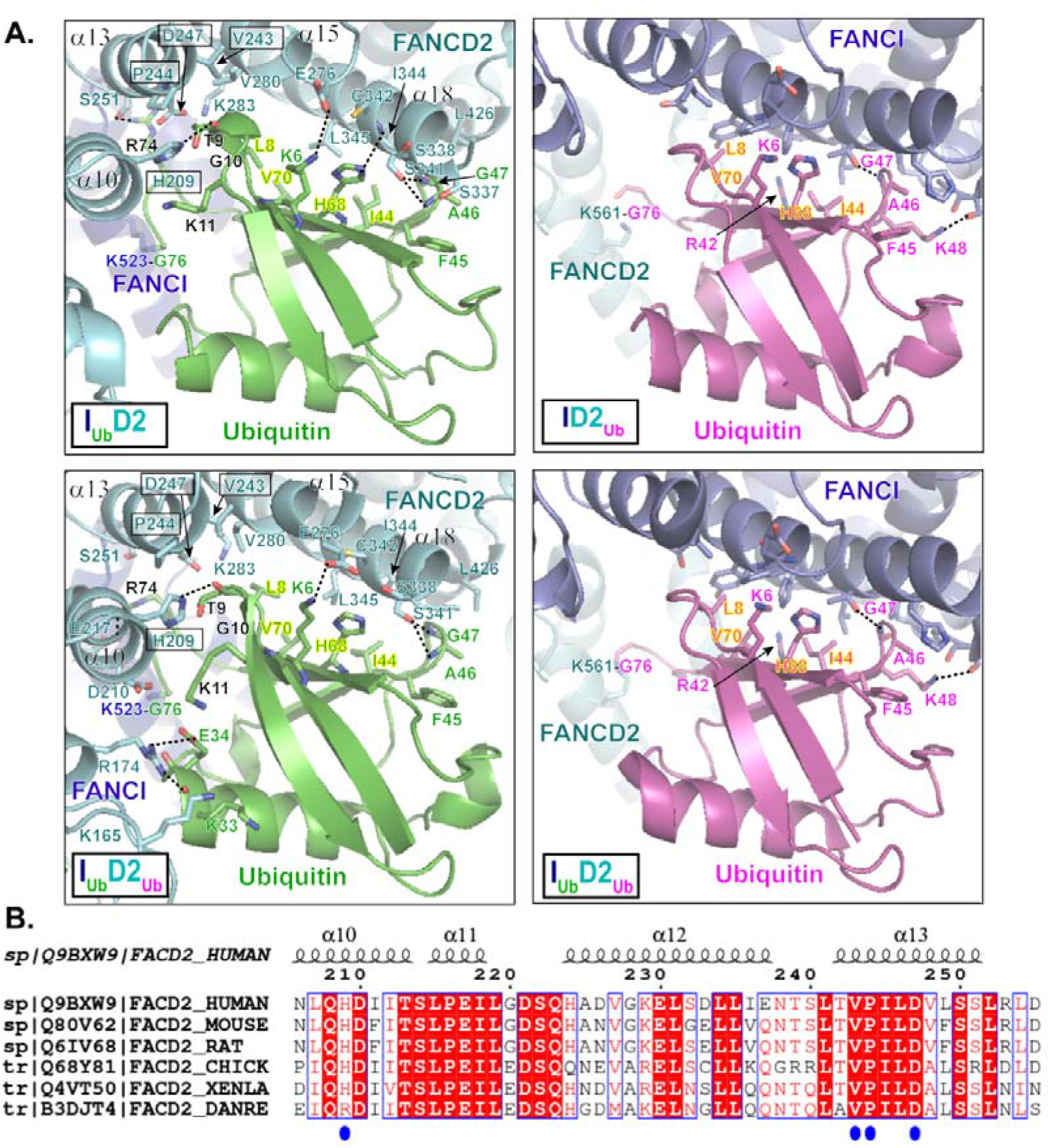
Interactions of FANCI/FANCD2 with the ubiquitin conjugated to the other ID2 subunit. **A**. Interactions of FANCI and FANCD2 with the ubiquitin conjugated to the other ID2 subunit, in DNA- bound I_Ub_D2, I_Ub_D2_Ub_ (PDB: 6VAE), ID2_Ub_ (PDB: 6VAF) and I_Ub_D2_Ub_ (PDB: 6VAE) structures (Wang *et al*, 2020). Dotted straight lines indicate hydrogen bonding. Both FANCD2 and FANCI interact with ubiquitin’s hydrophobic I44 patch (residues L8, I44, H68 and V70; all labelled in highlighted-yellow font) and additionally with residues F45 to G47 of ubiquitin. However, the ubiquitin conjugated to FANCI has a more extensive interface with FANCD2. This extended interface is formed by interactions of FANCD2 α10 - α13 helices (predominant interacting residues highlighted in boxes) with residues R74 and T9 to K11 of ubiquitin (shown in black font). The ubiquitin-FANCD2 interface may be further extended via interactions between residues K33 and E34 of ubiquitin with K165 and R174 of FANCD2, as shown in I_Ub_D2_Ub_-DNA structure. For direct comparison of corresponding interactions, the same orientation for all ubiquitins (both FANCI-conjugated and FANCD2- conjugated) was achieved by aligning: ID2_Ub_-DNA and I_Ub_D2_Ub_-DNA structures to the ubiquitin of I_Ub_D2-DNA structure, and subsequently, the I_Ub_D2_Ub_-DNA structure to the ubiquitin of ID2_Ub_-DNA structure as well. **B**. Clustal O multiple sequence alignment of human, mouse, rat, chicken, frog and zebrafish FANCD2 amino-acid sequences, focused on a region encompassing α11-13 helices of FANCD2 in human I_Ub_D2- DNA structure (helical regions shown on top). Identical residues among various species are highlighted red, whereas residues in positions displaying 83% similarity/identity are shown in red font. Residues of FANCD2 interacting with FANCI’s ubiquitin in both I_Ub_D2-DNA and I_Ub_D2_Ub_-DNA structures are indicated with blue circles.

## Discussion

ICLs and/or replication stress result in FA-core catalysed ID2 ubiquitination, which enables the ID2 complex to clamp on DNA (Lemonidis *et al*, 2021). Since ubiquitinated ID2 is able to slide on DNA *in vitro*, it has been proposed that ID2 ubiquitination effectively functions in sliding the ID2 complex away from ICLs/replication forks. This would allow nucleases or other factors to act for the repair of ICLs and/or restoration of replication, while the ID2 clamp may protect the DNA or have a processivity function (Wang *et al*, 2020). Loss of FANCD2 ubiquitination has been found to be equally bad for cell survival as loss of FANCD2, in response to the ICL-inducing agent, mitomycin C (Garcia-Higuera *et al*, 2001). In contrast, loss of FANCI ubiquitination has been shown to be less severe than loss of FANCI in similar cell-survival assays (Smogorzewska *et al*, 2007). Moreover, *in vivo* data show that blocking FANCD2 ubiquitination (K561R mutant) completely abolishes FANCI ubiquitination, whereas blocking FANCI ubiquitination (K523R mutant) greatly impairs, but does not completely abolish FANCD2 ubiquitination (Smogorzewska *et al*, 2007). Lastly, recent structural and biochemical evidence reveal that the FA-core complex and UBE2T preferentially target for ubiquitination the FANCD2 subunit of the ID2 complex, while FANCI ubiquitination lags (Wang *et al*, 2021).

The above indicate that in cells, FANCD2 ubiquitination most likely precedes FANCI ubiquitination, and because the two ubiquitination events are linked, FANCI ubiquitination is absolutely dependant on FANCD2 ubiquitination. Upon FANCD2 ubiquitination, the C-termini of FANCI and FANCD2 close around DNA, and this movement is associated with exposure of FANCI’s target lysine (K523) (Wang *et al*, 2020; Lemonidis *et al*, 2021). As a result, FANCI ubiquitination is greatly enhanced (Fig. 3). Indeed, and in agreement with what has been observed before with FA-core catalysed reactions (Wang *et al*, 2021), we found that the rate of FANCI ubiquitination is significantly higher in ID2_Ub_-DNA complex than in ID2-DNA complex (Fig. 3C). In the presence of DNA, FANCI ubiquitination is required for protecting FANCD2’s ubiquitin from excessive deubiquitination (Fig. 4A) (Arkinson *et al*, 2018; Rennie *et al*, 2020). Albeit slower in I_Ub_D2_Ub_-DNA complex, FANCD2 deubiquitination can still progress at significantly faster rate than FANCI deubiquitination (Fig. 1). Removal of FANCD2’s ubiquitin from the I_Ub_D2_Ub_-DNA complex does not impact on either the closed- on-DNA ID2 conformation (Fig. 2), or the high level protection of FANCI’s ubiquitin from USP1-UAF1- mediated deubiquitination. In fact, our deubiquitination assays indicate that I_Ub_ sensitivity to USP1- UAF1 action is conferred only by the absence of DNA, while enhanced protection is achieved when both FANCD2 and DNA are present (Fig. 4A). FANCD2’s target lysine (K561) is exposed for re-ubiquitination in the I_Ub_D2-DNA complex (Fig. 3A-B), and indeed the rate of FANCD2 ubiquitination in that complex is significantly greater than in the ID2-DNA complex (Fig. 3C). Hence, we propose a model whereby the balance between FANCD2 ubiquitination/deubiquitination determines whether FANCI gets ubiquitinated. Once FANCI ubiquitination is established, it plays a two-fold role: it prevents excessive FANCD2 deubiquitination (in I_Ub_D2_Ub_-DNA complex), and it ensures FANCD2 re-ubiquitination (in I_Ub_D2-DNA complex), once ubiquitin has been removed from FANCD2 (Fig. 6). The clamping on DNA of I_Ub_D2 and I_Ub_D2_Ub_ complexes ensures that maximum protection against USP1- UAF1 activity is achieved for both conjugated ubiquitins, and therefore ubiquitinated ID2 cannot easily revert to a non-ubiquitinated state. In essence, FANCI ubiquitination, via maintaining FANCD2 ubiquitination, commits the ID2_Ub_ complex for FA repair, since without FANCI ubiquitination such complex would be rapidly transformed to a non-ubiquitinated ID2 complex, through the action of USP1-UAF1.

**Fig. 6.**
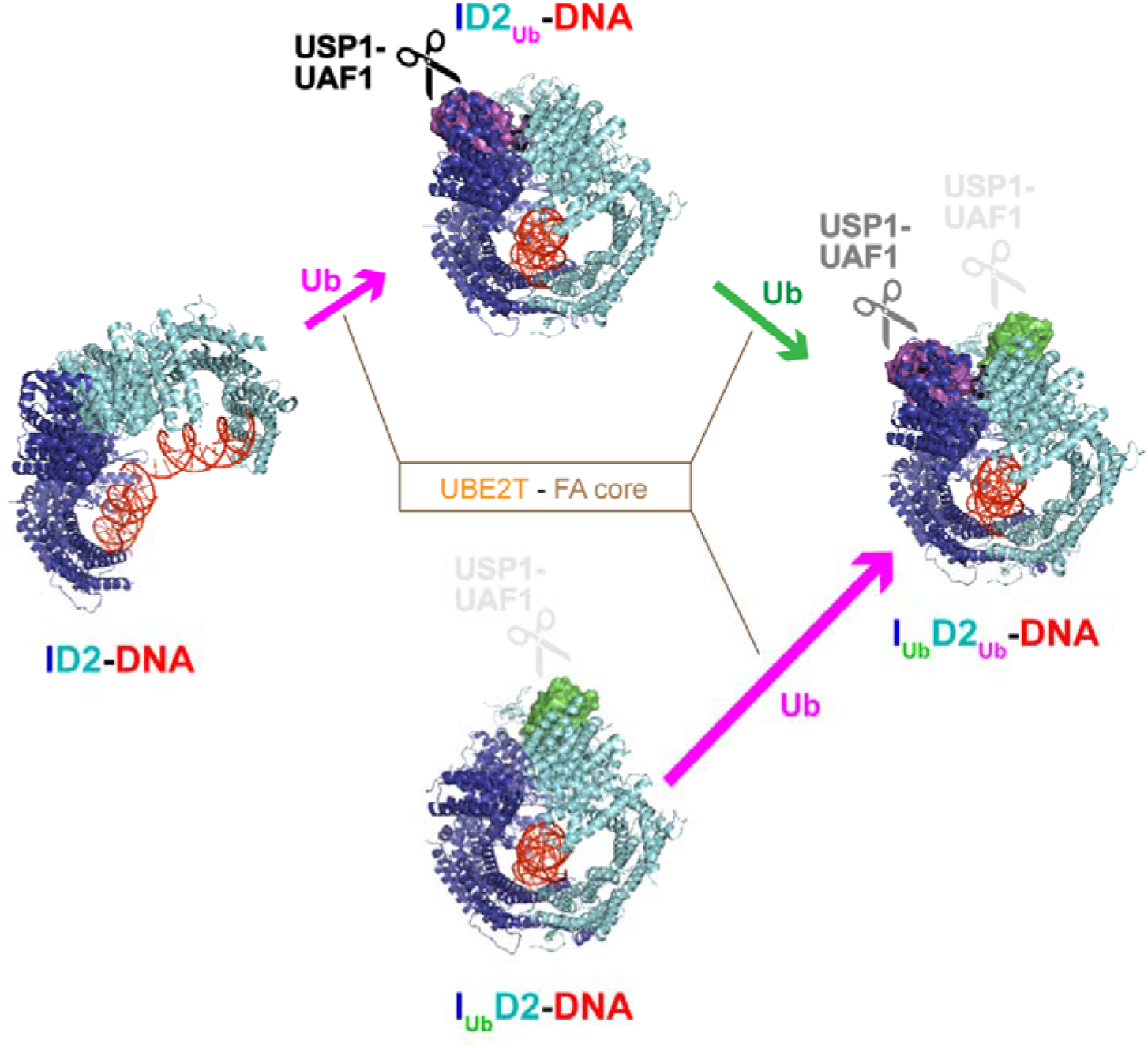
FANCI ubiquitination supports and maintains a di-mono-ubiquitinated ID2 state. Model explaining how the di-monoubiquitinated ID2 complex is generated and maintained. The UBE2T ubiquitin conjugating enzyme partners with the FA core ubiquitin ligase for ubiquitination of the DNA-bound FANCI-FANCD2 (ID2) complex. Of the two proteins subunits of the ID2 complex, FANCD2 is preferentially targeted for ubiquitination. While the resulting complex (ID2_Ub_-DNA) is sensitive to USP1-UAF1 deubiquitination activity, it has a conformation that now favours FANCI ubiquitination. Upon FANCI ubiquitination, the ubiquitin conjugated to FANCD2 gains some degree of resistance towards USP1-UAF1-mediated deubiquitination (I_Ub_D2_Ub_-DNA complex). Nevertheless, FANCD2’s ubiquitin is preferentially targeted for deubiquitination in the I_Ub_D2_Ub_-DNA complex. Its removal, though, is counteracted by very fast rates of FANCD2 ubiquitination (in the I_Ub_D2-DNA complex), which can (re-)establish the di-mono-ubiquitinated state (I_Ub_D2_Ub_-DNA). Since the ubiquitin-on-FANCI is highly protected against deubiquitination in both I_Ub_D2-DNA and I_Ub_D2_Ub_-DNA complexes, reverting to a non-ubiquitinated ID2 state is highly disfavoured, once FANCI ubiquitination is established. Arrow lengths are proportional to ubiquitination rates estimated in Fig 3C. ID2-DNA, ID2_Ub_-DNA and I_Ub_D2_Ub_-DNA structures shown correspond to PDB entries 6VAA, 6VAF and 6VAE, respectively (Wang *et al*, 2020).

It is worth noting that our ubiquitination reactions were performed using a minimal E1-E2-E3 system, consisting of a FANCL truncation mutant as a source of E3 ubiquitin ligase (FANCL^109-375^, instead of FA-core complex), and an engineered UBE2T (UBE2Tv4, displaying enhanced activity over wild type UBE2T), as source of E2 (Chaugule *et al*, 2019b, 2020). Nevertheless, we expect that, also under physiological conditions (ubiquitination with wild-type UBE2T and FA-core complex), FANCD2 would be ubiquitinated much faster within I_Ub_D2-DNA than within ID2-DNA complex, for the following reasons. Structural insights into ID2 ubiquitination by the FA-core complex, indicate that the FA-core is able to remodel the DNA-bound ID2 complex into a closed state conformation, whereby FANCD2’s target lysine and neighbouring residues are optimally positioned for ubiquitin-conjugation by the FANCL-bound UBE2T enzyme (Wang *et al*, 2021). Since I_Ub_D2-DNA is already in the closed state conformation, there is no need for the ID2 remodelling step to achieve FANCD2 ubiquitination. Such step is likely rate-limiting in FA-core catalysed ID2-DNA ubiquitination, since, both open-state and intermediated state ID2 conformations were additionally identified (and were equally distributed) in FA-core bound ID2 complexes, produced from such reactions (Wang *et al*, 2021). Hence, the reaction is expected to progress at a faster rate in the closed state I_Ub_D2-DNA than in the open-state ID2-DNA. Similarly, the rate of FANCI ubiquitination is significantly faster in the closed state ID2_Ub_-DNA complex than in the open state ID2-DNA complex, whether ubiquitination occurs utilizing the UBE2Tv4/FANCL^109-375^ pair (Fig 3C), or under more physiological (UBE2T/FA-core pair) conditions (Wang *et al*, 2021). Lastly, FANCD2 deubiquitination, which also involves a conformational transition step, whereby ID2_Ub_-DNA complex opens-up a bit upon USP1-UAF1 binding (Rennie *et al*, 2021), also progresses much faster when the initial closed ID2_Ub_-DNA state is compromised into a more open-state via the FANCI R1285A mutation (Wang *et al*, 2020).

Our results indicate that dsDNA, along with FANCI ubiquitination are required for maintaining FANCD2 ubiquitination, while dsDNA also protects from FANCI deubiquitination. This is in agreement with previous reports highlighting the protective role of DNA against FANCI/FANCD2 deubiquitination (van Twest *et al*, 2017; Arkinson *et al*, 2018). Interestingly, the opposite effect (DNA promoting USP1-UAF1-mediated FANCD2 deubiquitination) has been reported in a study utilizing a ∼60% FANCD2-ubiquitinated ID2 complex produced with the aid of a 64-mer single-stranded DNA (Liang *et al*, 2019). While the reasons for such discrepancy need to be further investigated, it is likely that the source of DNA (singe- versus double- stranded) used for ID2 ubiquitination, determines whether FANCD2 deubiquitination will be promoted or inhibited. UAF1 has been shown to bind both double-stranded (dsDNA), single-stranded (ssDNA), or more complex D-loop structures of DNA, *in vitro* (Liang *et al*, 2016, 2019). Although we cannot exclude the possibility of a UAF1-ssDNA binding event promoting ID2 deubiquitination, there is no evidence in support of dsDNA-UAF1 binding influencing ID2 deubiquitination: cryo-EM analysis of *in vitro* assembled USP1-UAF1-ID2_Ub_-DNA complexes, suggests that formation of a ternary complex is favoured, with both dsDNA and USP1- UAF1 preferentially binding ubiquitinated ID2, rather than each other (Rennie *et al*, 2021). Since dsDNA has been shown to protect against both FANCI and FANCD2 deubiquitination, ubiquitinated ID2 complexes may need to be disengaged from DNA to be more effectively deubiquitinated. This could be achieved through the action of the DVC1-p97 ubiquitin segregase, which has been shown to be responsible for removal of ID2 from sites of DNA damage, once ID2 has been SUMOylated and subsequently polyubiquitinated on SUMO (Gibbs-Seymour *et al*, 2015). Another possibility would be that ubiquitinated ID2 and/or USP1/UAF1 are modulated (by factors and in ways that are yet unknown) for effective cleavage of the conjugated ubiquitins, in the presence of DNA.

The exact mechanism by which dsDNA is protecting both FANCD2 and FANCI from deubiquitination is yet unclear. For the protection seen on I_Ub_ in the absence of D2, there is a possibility that dsDNA blocks access of USP1-UAF1 to FANCI’s ubiquitin, either directly or indirectly through altering the conformation of I_Ub_. Further structural work will be required to elucidate how USP1-UAF1 targets the ubiquitin on FANCI and how dsDNA may interfere with such targeting. The deubiquitination protection of I_Ub_D2 and I_Ub_D2_Ub_ complexes by dsDNA, however, is likely mediated through stabilisation of the ubiquitinated ID2 complexes in the closed conformation in which FANCI’s ubiquitin, and therefore FANCD2’s ubiquitin too, are maximally protected. In support for this, the R1285Q mutant of FANCI, which is predicted to disrupt the closed-on-DNA conformation of ubiquitinated ID2 complexes (Wang *et al*, 2020; Rennie *et al*, 2020)(Fig. EV1), impairs both the dsDNA binding (Rennie *et al*, 2020) and the I_Ub_/D2_Ub_ protection from USP1-UAF1-mediated deubiquitination (Wang *et al*, 2020). While closed-conformation ubiquitinated ID2 complexes can exist in the absence of DNA, as previously shown for ID2_Ub_ (Rennie *et al*, 2020), such DNA-free complexes may be less stable, or conformationally more flexible, without the avidity conferred by the interacting DNA. Therefore, they may be more amenable to deubiquitination. Our biochemical assays indicate that FANCI ubiquitination further secures this closed ID2 conformation. This is likely achieved through FANCI-ubiquitination effectively restricting the conformational inclination of the USP1-UAF1-bound ID2 complex towards the open-state conformation. Indeed, the structure of USP1-UAF1 complex bound to ID2_Ub_-DNA revealed ID2 movements towards the open-state conformation, affecting not only FANCI helices in the region where ubiquitin-conjugation occurs, but also, and most profoundly, the FANCD2 N-terminus involved in interaction with FANCI’s ubiquitin (Rennie *et al*, 2021).

In ubiquitinated ID2 complexes, the ubiquitin conjugated to FANCI is substantially more protected from deubiquitination than the ubiquitin conjugated on FANCD2. To some extent, this may be due to USP1-UAF1 preferentially targeting FANCD2, via USP1’s N-terminal extension (Arkinson *et al*, 2018) and through UAF1-FANCI interactions acting as a USP1-FANCD2 enzyme-substrate recruitment module (Rennie *et al*, 2021). Nevertheless, the preferential targeting of FANCD2’s ubiquitin over FANCI’s ubiquitin may also be due to the latter ubiquitin participating in more extensive interactions (than the former ubiquitin) with the other ID2 subunit (Fig. 5A)(Wang *et al*, 2020; Rennie *et al*, 2021). Further biochemical work, including extensive mutagenesis of key ubiquitin-ID2 interfaces, is likely to uncover the reasons contributing to D2_Ub_ being preferentially targeted over I_Ub_ in ubiquitinated ID2 complexes.

In this work we provide a structural and biochemical basis for the *in vivo* interdependency in FANCI and FANCD2 ubiquitination observed before (Smogorzewska *et al*, 2007; Sims *et al*, 2007). This is crucial for understanding how FANCI and FANCD2 ubiquitination/deubiquitination are linked at the molecular level. However, while the mechanism of FANCD2 ubiquitination and deubiquitination has been sufficiently elucidated (Chaugule *et al*, 2020; Wang *et al*, 2021; Rennie *et al*, 2021), we are still lacking information on how UBE2T and FA-core engage the mono-ubiquitinated ID2 complex for FANCI ubiquitination, and how the ubiquitin from FANCI is removed by USP1-UAF1. Deciphering how FANCI ubiquitination and deubiquitination are encoded as well, coupled with generation of mutants affecting FANCI-only and/or FANCD2-only ubiquitination/deubiquitination, would allow us to study in more detail how these processes are dynamically regulated *in vivo*.

## Methods

### Protein expression and purification

Protein constructs for protein expression were as before (Arkinson *et al*, 2018; Rennie *et al*, 2020). All proteins and ubiquitinated versions of FANCI and FANCD2 were produced as previously described, in the absence of DNA, and ID2 complexes (with or without DNA) were subsequently assembled *in vitro* (Arkinson *et al*, 2018; Rennie *et al*, 2020). Briefly, FANCI, FANCD2, USP1 and USP1ΔN proteins, corresponding to canonical human protein sequences, were expressed with N-terminal six-histidine tag fusions in Sf21 insect cells and were subsequently purified using NiNTA chromatography, anion exchange and gel filtration. Untagged human UAF1 was co-expressed and co-purified with USP1, whereas his-tagged UAF1 was expressed and purified in isolation, to be later used for *in vitro* assembly of USP1ΔN-UAF1 complex. For production of ubiquitinated FANCI and FANCD2 proteins, reactions occurred using FANCI/FANCD2, UBA1, UBE2T (UBE2Tv4), FANCL^109-375^, ATP-Mg^+2^ and, either Spy-tagged ubiquitin, or GST-tagged ubiquitin (both tags were N-terminal). In the case of Spy-tagged ubiquitin reactions, incubation with GST-tagged SpyCatcher occurred afterwards to covalently link GST to ubiquitin. Ubiquitinated proteins were then purified by capture of GST-linked ubiquitin on Glutathione resin, and subsequent gel filtration of ubiquitinated products. After final gel filtration step, in buffer containing 20 mM Tris pH8, 100 mM NaCl (or 400 mM for ubiquitinated/non-ubiquitinated FANCI/FANCD2), 5% glycerol and reducing agent (0.5-1 mM TCEP or 2-5 mM DTT), proteins were flash frozen in liquid nitrogen and stored at -80° C. Ubiquitinated (ID2_Ub_, I_Ub_D2 and I_Ub_D2_Ub_) or non-ubiquitinated ID2 complexes were assembled on ice from individually purified proteins equilibrated in gel-filtration buffer having 150 mM NaCl concentration.

### DNA oligos

All DNA oligos were purchased from IDT and consisted of perfectly complementary pairs for formation of double stranded DNA (dsDNA) molecules. Unlabelled DNA oligos (for 32 bp, 51 bp and 61 bp dsDNA formation) were PAGE-purified, whereas 5’-labelled with IRDye700 oligos (for infrared-labelled 32 bp dsDNA formation) were HPLC purified. Their 5’ to 3’ sequence is as following: 32 bp (labelled/unlabelled): CGATCGGTAACGTATGCTGAATCTGGTGCTGG and corresponding complementary sequence; 51 bp: CGTCGACTCTACATGAAGCTCGAAGCCATGAATTCAAATGACCTCTGATCA and corresponding complementary sequence; and 61 bp: TGATCAGAGGTCATTTGAATTCATGGCTTCGAGCTTCATGTAGAGTCGACGGTGCTGGGAT and corresponding complementary sequence.

### Cryo-EM sample preparation, data collection and processing

Purified ubiquitinated His_6_-V5-FANCI was mixed with purified FANCD2 at 1:1 molar ratio and exchanged into cryo-EM buffer (20 mM Tris, pH 8.0, 150 mM NaCl, 2 mM DTT) using a Bio-Spin P-30 column (Bio-Rad). The concentration of the recovered protein complex was estimated based on its absorbance at 280 nM. A PAGE-purified 61 base-pair dsDNA was then added to the protein complex at a 1:1 molar ratio. After a short equilibration at room temperature, 3.5 µl of the protein-DNA mix (7.6 µM) was loaded on Quantifoil 1.2/1.3 300 mesh grids, which had been previously glow discharged for 30 sec at 30 mA. Grids were blotted for 3 sec and vitrified in liquid ethane using a Vitrobot operating at 95% humidity at 18 °C. The frozen grids were subsequently imaged over two sessions on CRYO ARM 300 (JEOL) microscope (Scottish Centre for Macromolecular Imaging) using a DE64 detector. For the second session, the in-column omega filter was used, with a slit width of 30 eV. 45-frame movies (11,229 in total), with a calibrated pixel size of 1.023 Å, were collected in counting mode, using serialEM software (Mastronarde, 2005). Total electron dose was either 45.2 e/Å^2^ or 46.8 e/Å^2^ over 15.32 seconds. Movies were subsequently processed in cryoSPARC (v3.2) (Punjani *et al*, 2017) for particle-image extraction, 2D classification and construction/refinement of cryo-EM density maps, as detailed below. Each set of movies (from the two sessions) was processed separately for obtaining good particles. Following Patch motion correction, Patch CTF estimation and curation of resulting exposures, we obtained 10,178 dose-weighted motion-corrected images in total. Particles were first picked automatically using elliptical blobs having minimum and maximum diameters of 120 and 200 Å, respectively. All picked particles were extracted within a 320-pixel box. Following few rounds of 2D classification, particles forming good 2D classes were used for ab-initio 3D reconstruction (2-3 models) followed by heterogeneous refinement. To clear junk particles, the initial 3D classes were subjected to further rounds of heterogeneous refinement, using each time as an input particle-set the good 3D class output-particles of the previous hetero-refinement job. We then subjected the particles of the good 3D class to 2D classification to generate 2D templates for particle picking. The particle maximum diameter was set to 200 Å, in template-based particle picking. Template picking occurred twice with a different set of templates each time for both micrograph datasets. Removal of junk particles occurred with heterogeneous refinement, as before: all template-based extracted particle picks were subjected to multiple rounds of heterogeneous refinement, using one previously generated good, and 1-2 previously generated junk 3D classes. Multiple rounds of heterogeneous refinement was also applied to all particles extracted by blob-picking to enrich the good particle set. To further enrich the good particles of some 3D classes, ab-initio reconstruction (2-3 models) followed by heterogeneous refinement also occurred for these classes. After removal of any duplicate particles among the eight generated 3D classes (4 for each dataset) and another round of heterogeneous refinement, the 259,775 particles falling to the good 3D class were motion-corrected locally and re-extracted from the micrographs. Following another round of heterogeneous refinement, the resulting good 3D class (made of 206,669 particles) was low-pass filtered at 12, 18 and 30 Å. A final round of heterogeneous refinement then occurred using the three low passed filtered volumes and a starting refinement resolution set at 12 Å. The resulting highest resolution class (made of 139,601 particles) resulting from the 12 Å filtered map, was both homogeneous and non-uniform refined. We then applied a local non-uniform refinement (Punjani *et al*, 2020), using as inputs, the output volume of the non-uniform refinement job, and the mask generated by the homogeneous refinement job. The generated map had an overall resolution of 4.14 Å, determined by gold-standard FSC. A local resolution filtered map was then obtained, by calculating the local resolution at 0.143 FSC threshold with an adaptive window factor of eight. This map was used for model building, while a sharpened map produced with Phenix (v1.19.2) auto- sharpen tool (Terwilliger *et al*, 2018) was also used to aide interpretation of higher resolution features.

### Model building and visualization

Initially, the ubiquitin conjugated to FANCD2 in the I_Ub_D2_Ub_-DNA structure (PDB code: 6VAE)(Wang *et al*, 2020) was removed, and the remaining structure was fitted to the I_Ub_D2-DNA map using Chimera software. Model editing and building subsequently occurred in WinCoot (v0.9.4.1) (Emsley *et al*, 2010), incorporating torsion, planar peptide, trans peptide and Ramachandran restraints. More specifically: i) we corrected for peptide twists and mismatches to FANCI and FANCD2 human protein sequences (UniProt entries: Q9NVI1 and Q9BXW9, respectively); ii) we removed regions corresponding to poor density - such as a large section of FANCI N-terminus (residues 1-171), few FANCI/FANCD2 loops and a short stretch in one end (2 bp) of the 29bp dsDNA (closer to FANCD2 C-terminus); and iii) we filled some gaps in the structural model for which the cryo-EM density was sufficiently good for modelling building. Then, we performed several rounds of, (global and local) automated real-space refinement in Phenix (v1.19.2) (Afonine *et al*, 2018), followed by manual refinement of problematic regions/residues in WinCoot. For automated refinement, protein/dsDNA secondary structure, rotamer, Ramachadran, geometry and FANCI-Ub K523-G76 bond restrains were enforced. Additionally, we applied a non-bonded weight of 2500 to restrict steric clashes. Cryo-EM data and model refinement statistics are reported in Table 1. Maps and models were visualized in PyMOL (The PyMOL Molecular Graphics System, Version 1.7.6.6 Schrödinger, LLC.), Chimera (Pettersen *et al*, 2004), or ChimeraX (Goddard *et al*, 2018). Surface accessibility of non-conjugated FANCI (K523) and FANCD2 (K561) target lysines in our I_Ub_D2-DNA structure, and in ID2-DNA (PDB:6VAA) and ID2_Ub_ (PDB:6VAF) structures (Wang *et al*, 2020), was measured using the PDBePISA tool (Krissinel & Henrick, 2007) at https://www.ebi.ac.uk/pdbe/pisa/.

### In vitro reactions

FANCI-FANCD2 reactions occurred in 10 µl volume, using FLAG-tagged FANCD2 and/or His- V5-tagged FANCI (Both N-terminally tagged). *In vitro* ubiquitination reactions were conducted at 30° C with UBA1 (50 nM), UBE2Tv4 (2 µM), FANCL^109-375^ (2 µM), ubiquitin (10 µM), ID2 (2 µM) and a 32 bp dsDNA (3.6 µM), in 42 mM Tris-HCl pH 8, 140 mM NaCl, 5% Glycerol, 5 mM ATP, 5 mM MgCl_2_ and 1mM DTT. *In vitro* deubiquitination reactions occurred at room temperature, with USP1-UAF1 (50 nM) and ubiquitinated FANCI/ID2 (0.5 µM), in 50 mM Tris pH8, 100 mM NaCl, 5% Glycerol, 2 mM DTT. Unless otherwise stated, deubiquitination reactions were performed in the presence of 2 µM dsDNA (51 bp). Ubiquitination/deubiquitination reactions were terminated by addition of reducing LDS sample buffer (to 1x final). After boiling for 3 min at 100° C, a fraction of these (amount corresponding to approximately 0.5 pmoles of total ID2) were loaded on 4-12% Wedge-Well Tris-Glycine gels (Thermo Fisher). Following SDS-PAGE, proteins were transferred to nitrocellulose membranes, using an iBlot gel transfer device (Thermo Fisher). FLAG-FANCD2 and His_6_-V5-FANCI were visualized, on Odyssey CLx (LI-COR) infrared scanner, following western blotting with FANCD2 (sc-28194; Santa Cruz Biotechnology) and V5 (66007.1-Ig; ProteinTech) primary antibodies, and corresponding infrared-dye-conjugated secondary antibodies, as described before (Rennie *et al*, 2020).

### Protein Induced Fluorescence Enhancement assays

These were performed using ubiquitinated or non-ubiquitinated His_6_-V5-FANCI, non-ubiquitinated His_6_-FANCD2 and infrared (IRDye-700) labelled (on both strands) 32 bp DNA in Fluorescence Buffer (20 mM Tris pH 8.0, 150 mM NaCl, 5% glycerol, 0.47 mg/ml BSA, 1 mM DTT). PIFE assays were conducted as before (Rennie *et al*, 2020), but with the following modifications: Ubiquitinated/non-ubiquitinated His_6_-V5-FANCI was first diluted to 5 µM and then subjected to several two-fold serial dilutions, whereas His_6_-FANCD2 was mixed with labelled DNA to achieve a working concentration of 5 µM FANCD2 and 250 nM DNA. Then 5 µl of this FANCD2-DNA mix was mixed with 5 µl of each of the FANCI series of dilutions for final concentrations of: 125 nM DNA, 2.5 µM FANCD2 and 1.2 nM -2.5 µM of FANCI in Fluorescence Buffer. For assessing FANCD2’s affinity to dsDNA, His_6_-FANCD2 was subjected to several two-fold serial dilutions and then each of this was mixed with a constant concentration of labelled DNA to achieve final concentration of 125 nM DNA and 40 nM - 5 µM of FANCD2 in Fluorescence Buffer. Samples to be measured were transferred into premium capillaries (NanoTemper Technologies). Measurements were performed at 22°C on a Monolith NT.115 instrument (NanoTemper Technologies) using the red channel, with laser power set to 20%.

### Quantification and statistical analysis

For ubiquitination/deubiquitination experiments, the percentage FANCI/FANCD2 ubiquitination (induced-by-ubiquitination or residual-from-deubiquitination) at indicative time- points, was calculated following quantification of ubiquitinated and non-ubiquitinated FANCI/FANCD2 bands from western blots, using LI-COR Image Studio Lite software (v5.2). All the percentage ubiquitination values calculated for each complex/protein from multiple experiments were used in fitting to either a one phase decay (deubiquitination experiments) or a one-phase association (ubiquitination experiments) model, assuming same plateau for all proteins analysed. Assessment of statistically significant changes was done using one-way ANOVA with Bonferroni correction. Dissociation constants with associated uncertainties from Protein-Induced Fluorescence Enhancement (PIFE) assays were determined by fitting baseline subtracted PIFE values to a one-site binding model, as described before (Rennie *et al*, 2020). All data deriving from quantification of blots and PIFE experiments were visualized and statistically analysed using GraphPad Prism software.

## Data availability

The two half maps of the final cryo-EM I_Ub_D2-DNA complex reconstruction, along with the locally filtered and Phenix Autosharpen maps deriving from these, have been deposited to the Electron Microscopy Data Bank with accession code EMD-14694. The atomic coordinates of the refined model have been deposited to the Protein Data Bank with accession code 7ZF1.

## Author contribution

KL conceived, designed and performed the experiments, wrote the manuscript and made the figures. MLR conceived and designed PIFE experiments which he executed together with KL. MLR also helped KL with cryo-EM processing and structure modelling. KL, MLR, CA and VKC expressed and purified proteins. MC prepared grids. JS collected cryo-EM data. HW secured funding for the project. KL edited the manuscript with help from MLR, CA, JS and HW. All authors approved the final version of the manuscript.

## Acknowledgments

Access to cryo-EM instrumentation was provided by the Scottish Centre for Macromolecular Imaging (SCMI), funded by the MRC (MC_PC_17135) and SFC (H17007). This work was supported by the European Research Council (ERC- 2015-CoG-681582 ICLUb) consolidator grant to HW.

## Tables and Figures

**Fig. EV1.**
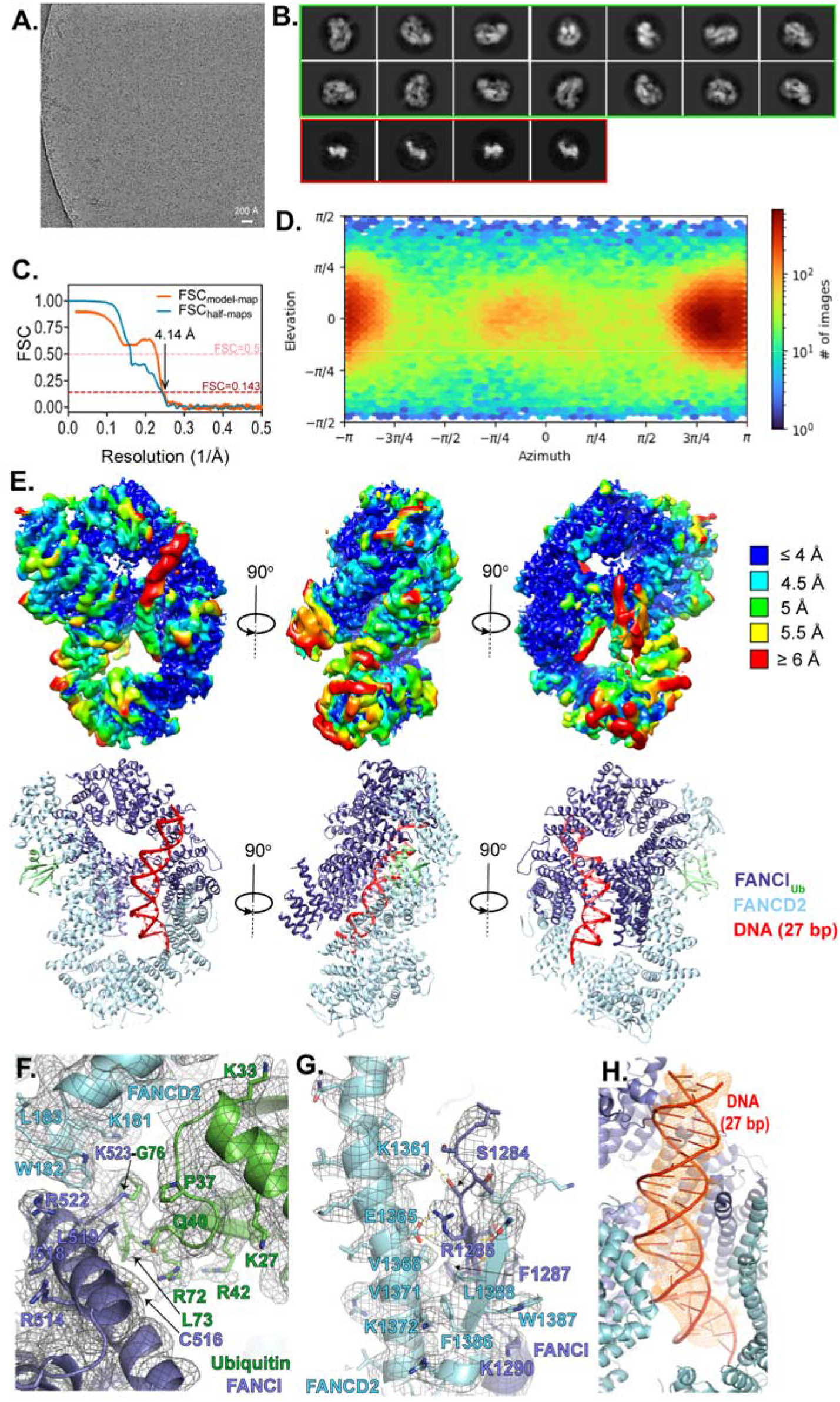
Cryo-EM analysis and structure modelling of I_Ub_D2-DNA complex. **A**. Example micrograph with scale bar. **B**. Example 2D classes. Circular mask is 170Å in diameter. 2D classes surrounded by a green box correspond to I_Ub_D2-DNA complex particles, while smaller-sized 2D classes surrounded by a red box likely correspond to monomeric I_Ub_/D2 proteins. **C**. Fourier Shell Correlation (FSC) curves: between the two half maps produced in the final local non- uniform refinement (shown in blue) and between the refined model and final map (shown in orange). **D**. Particle orientation (viewing direction distribution) in the final map. Total number of particles: 139,601. **E**. *Top*: Locally filtered map coloured by local resolution, at three different angles. *Bottom*: Corresponding structural model viewed under same orientations. **F**. I_Ub_D2-DNA structure with corresponding map density (locally filtered map), centred on the isopeptide bond between K523 of FANCI and G76 of ubiquitin. Some well-resolved side-chains are illustrated as sticks and indicated. **G**. Interaction between FANCI and FANCD2 C-termini with corresponding map density (locally filtered map). A beta-sheet consisting of a FANCI and a FANCD2 strand is formed (residues 1285- 1289 of FANCI and residues 1384-88 of FANCD2). This is held in place through hydrophobic and electrostatic interactions with a FANCD2 helix (1351-1377 aa). Residues predicted to participate in such interactions are shown as sticks and indicated. Selected side chains, for which there is good density are also shown as sticks. For clarity, adjacent to that region elements of the I_Ub_D2-DNA structure and map are removed. **H**. I_Ub_D2-DNA structure centred on DNA. Density corresponding to the 27 bp modelled DNA is shown as orange mesh. Colouring of structure is as in E-G.

**Fig. EV2.**
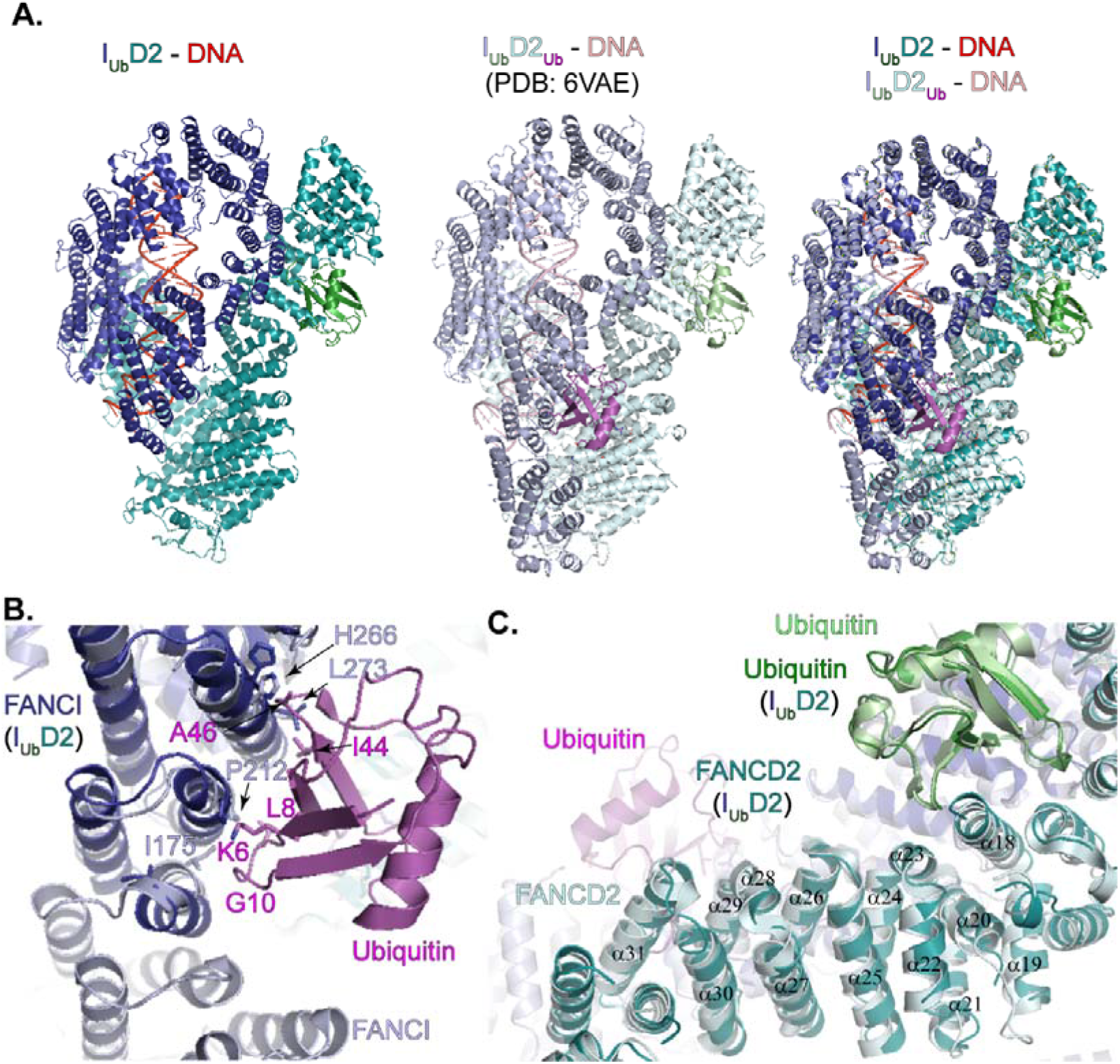
I_Ub_D2-DNA and I_Ub_D2_Ub_-DNA structure comparison. **A**. The lack of the FANCD2-conjugated ubiquitin in I_Ub_D2-DNA structure is associated with a disorder in the 170 aa N-terminus of FANCI, when compared with I_Ub_D2_Ub_-DNA structure. The two structures were aligned in Pymol and visualized from the same angle, either on their own (*left* and *centre*), or together (*right*). **B**. Helices and corresponding residues of FANCI involved in interaction with FANCD2’s ubiquitin (I_Ub_D2_Ub_-DNA structure), are positioned differently when the ubiquitin is removed (I_Ub_D2-DNA structure). Structures shown in A, were centred on FANCI N-terminus. Residues predominantly involved in ubiquitin-FANCI interactions are indicated and shown as sticks. The corresponding FANCI residues in I_Ub_D2-DNA structure are also shown as sticks. **C**. Removal of ubiquitin (magenta) from FANCD2 results in slight movements affecting several FANCD2 helices, from α31 (helix where ubiquitin is conjugated) up to α18. FANCD2 helices of I_Ub_D2- DNA and I_Ub_D2_Ub_-DNA are better aligned towards the N-terminus of FANCD2, where interaction with the ubiquitin (green) conjugated to FANCI occurs (N-terminally to, and including α18 helix of FANCD2). The structures shown in A, were centred on the central part of FANCD2.

**Fig. EV3.**
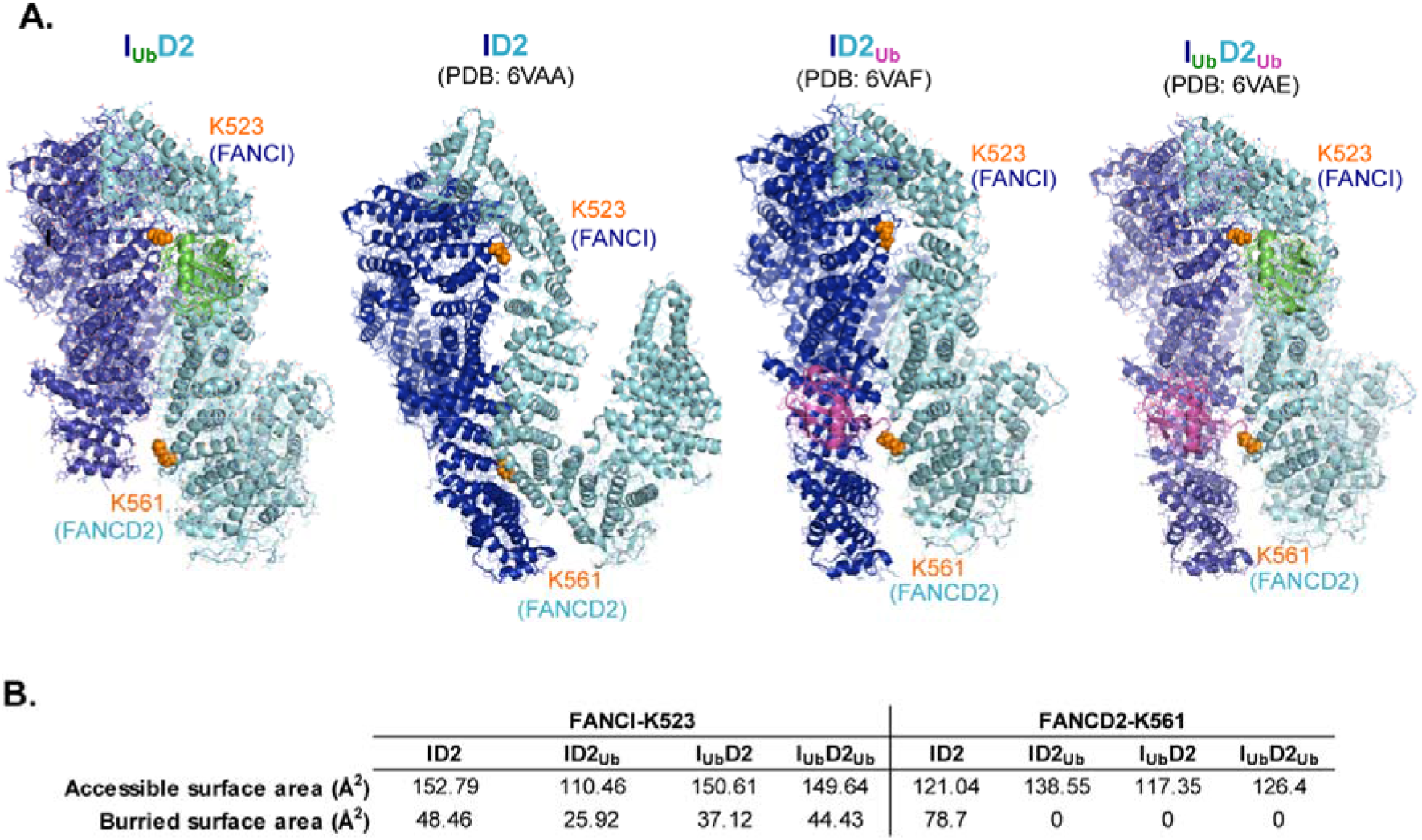
FANCI and FANCD2 target lysine positioning and accessibility in DNA-bound I_Ub_D2, ID2, ID2_Ub_ and I_Ub_D2_Ub_ structures. **A**. The overall accessibility of FANCI’s K523 and FANCD2’s K561 is shown within I_Ub_D2-DNA, ID2-DNA (PDB: 6VAA), ID2_Ub_-DNA (PDB: 6VAF) and I_Ub_D2_Ub_-DNA (PDB:6VAE) structures. For clarity DNA was removed from the structures. The corresponding lysines are illustrated as orange spheres. I_Ub_D2, ID2_Ub_ and I_Ub_D2_Ub_ structures were aligned to FANCI of ID2 structure, to allow visualization of all structures under same orientation. **B**. FANCI’s K523 and FANCD2’s K561 accessible surface areas and buried surface area (both in Å^2^) are shown for each of the structures illustrated in A. These values were determined from associated PDB files using the PDBePISA tool (Krissinel & Henrick, 2007) at https://www.ebi.ac.uk/pdbe/pisa/.

## References

Afonine P V., Poon BK, Read RJ, Sobolev O V., Terwilliger TC, Urzhumtsev A & Adams PD (2018) Real-space refinement in PHENIX for cryo-EM and crystallography. urn:issn:2059-7983 74: 531–544

Alcón P, Shakeel S, Chen ZA, Rappsilber J, Patel KJ & Passmore LA (2020) FANCD2–FANCI is a clamp stabilized on DNA by monoubiquitination of FANCD2 during DNA repair. Nat Struct Mol Biol 27: 240–248

Arkinson C, Chaugule VK, Toth R & Walden H (2018) Specificity for deubiquitination of monoubiquitinated FANCD2 is driven by the N-terminus of USP1. Life Sci Alliance 1: e201800162

Chaugule VK, Arkinson C, Rennie ML, Kamarainen O, Toth R & Walden H (2019a) Allosteric mechanism for site-specific ubiquitination of FANCD2.

Chaugule VK, Arkinson C, Rennie ML, Kämäräinen O, Toth R & Walden H (2020) Allosteric mechanism for site-specific ubiquitination of FANCD2. Nat Chem Biol 16: 291–301

Chaugule VK, Arkinson C, Toth R & Walden H (2019b) Enzymatic preparation of monoubiquitinated FANCD2 and FANCI proteins. Methods Enzymol 618: 1–30

Chen J, Dexheimer TS, Ai Y, Liang Q, Villamil MA, Inglese J, Maloney DJ, Jadhav A, Simeonov A & Zhuang Z (2011) Selective and Cell-Active Inhibitors of the USP1/ UAF1 Deubiquitinase Complex Reverse Cisplatin Resistance in Non-small Cell Lung Cancer Cells. Chem Biol 18: 1390–1400

Emsley P, Lohkamp B, Scott WG & Cowtan K (2010) Features and development of Coot. urn:issn:0907-4449 66: 486–501

Garcia-Higuera I, Taniguchi T, Ganesan S, Meyn MS, Timmers C, Hejna J, Grompe M & D’Andrea AD (2001) Interaction of the Fanconi anemia proteins and BRCA1 in a common pathway. Mol Cell 7: 249–262

García-Santisteban I, Peters GJ, Giovannetti E & Rodríguez JA (2013) USP1 deubiquitinase: Cellular functions, regulatory mechanisms and emerging potential as target in cancer therapy. Mol Cancer 12: 1–12

Gibbs-Seymour I, Oka Y, Rajendra E, Weinert BT, Passmore LA, Patel KJ, Olsen J V., Choudhary C, Bekker-Jensen S & Mailand N (2015) Ubiquitin-SUMO circuitry controls activated fanconi anemia ID complex dosage in response to DNA damage. Mol Cell 57: 150–164

Goddard TD, Huang CC, Meng EC, Pettersen EF, Couch GS, Morris JH & Ferrin TE (2018) UCSF ChimeraX: Meeting modern challenges in visualization and analysis. Protein Sci 27: 14–25

Kim JM, Parmar K, Huang M, Weinstock DM, Ruit CA, Kutok JL & D’Andrea AD (2009) Inactivation of Murine Usp1 Results in Genomic Instability and a Fanconi Anemia Phenotype. Dev Cell 16: 314–320

Krissinel E & Henrick K (2007) Inference of Macromolecular Assemblies from Crystalline State. J Mol Biol 372: 774–797

KSQ Therapeutics Inc (2021) A Phase 1 Study of KSQ-4279 Alone and in Combination in Patients With Advanced Solid Tumors. Identifier: NCT05240898: https://clinicaltrials.gov/ct2/show/NCT05240898

Lemonidis K, Arkinson C, Rennie ML & Walden H (2021) Mechanism, specificity, and function of FANCD2-FANCI ubiquitination and deubiquitination. FEBS J doi:10.1111/febs.16077 [PREPRINT]

Liang F, Longerich S, Miller AS, Tang C, Buzovetsky O, Xiong Y, Maranon DG, Wiese C, Kupfer GM & Sung P (2016) Promotion of RAD51-Mediated Homologous DNA Pairing by the RAD51AP1-UAF1 Complex. Cell Rep 15: 2118–2126

Liang F, Miller AS, Longerich S, Tang C, Maranon D, Williamson EA, Hromas R, Wiese C, Kupfer GM & Sung P (2019) DNA requirement in FANCD2 deubiquitination by USP1-UAF1-RAD51AP1 in the Fanconi anemia DNA damage response. Nat Commun 10: 2849

Liang Q, Dexheimer TS, Zhang P, Rosenthal AS, Villamil MA, You C, Zhang Q, Chen J, Ott CA, Sun H, et al (2014) A selective USP1-UAF1 inhibitor links deubiquitination to DNA damage responses. Nat Chem Biol 10: 298–304

Lim KS, Li H, Roberts EA, Gaudiano EF, Clairmont C, Sambel LA, Ponnienselvan K, Liu JC, Yang C, Kozono D, et al (2018) USP1 Is Required for Replication Fork Protection in BRCA1-Deficient Tumors. Mol Cell 72: 925–941.e4

Liu W, Palovcak A, Li F, Zafar A, Yuan F & Zhang Y (2020) Fanconi anemia pathway as a prospective target for cancer intervention. Cell Biosci 2020 101 10: 1–14

Longerich S, Kwon Y, Tsai MS, Hlaing AS, Kupfer GM & Sung P (2014) Regulation of FANCD2 and FANCI monoubiquitination by their interaction and by DNA. Nucleic Acids Res 42: 5657–5670

Ma A, Tang M, Zhang L, Wang B, Yang Z, Liu Y, Xu G, Wu L, Jing T, Xu X, et al (2018) USP1 inhibition destabilizes KPNA2 and suppresses breast cancer metastasis. Oncogene 2018 3813 38: 2405–2419

Ma L, Lin K, Chang G, Chen Y, Yue C, Guo Q, Zhang S, Jia Z, Huang TT, Zhou A, et al (2019) Aberrant activation of b-catenin signaling drives glioma tumorigenesis via USP1-mediated stabilization of EZH2. Cancer Res 79: 72–85

Mastronarde DN (2005) Automated electron microscope tomography using robust prediction of specimen movements. J Struct Biol 152: 36–51

Murai J, Yang K, Dejsuphong D, Hirota K, Takeda S & D’Andrea AD (2011) The USP1/UAF1 complex promotes double-strand break repair through homologous recombination. Mol Cell Biol 31: 2462–9

Mussell A, Shen H, Chen Y, Mastri M, Eng KH, Bshara W, Frangou C & Zhang J (2020) USP1 Regulates TAZ Protein Stability Through Ubiquitin Modifications in Breast Cancer. Cancers (Basel) 12: 3090

Nalepa G & Clapp DW (2018) Fanconi anaemia and cancer: an intricate relationship. Nat Rev Cancer 18: 168–185

Niraj J, Färkkilä A & D’Andrea AD (2019) The Fanconi Anemia Pathway in Cancer. Annu Rev Cancer Biol 3: 457–478

Niu Z, Li X, Feng S, Huang Q, Zhuang T, Yan C, Qian H, Ding Y, Zhu J & Xu W (2020) The deubiquitinating enzyme USP1 modulates ERα and modulates breast cancer progression. J Cancer 11: 6992–7000

Oestergaard VH, Langevin F, Kuiken HJ, Pace P, Niedzwiedz W, Simpson LJ, Ohzeki M, Takata M, Sale JE & Patel KJ (2007) Deubiquitination of FANCD2 Is Required for DNA Crosslink Repair. Mol Cell 28: 798–809

Pettersen EF, Goddard TD, Huang CC, Couch GS, Greenblatt DM, Meng EC & Ferrin TE (2004) UCSF Chimera--a visualization system for exploratory research and analysis. J Comput Chem 25: 1605–1612

Punjani A, Rubinstein JL, Fleet DJ & Brubaker MA (2017) CryoSPARC: Algorithms for rapid unsupervised cryo-EM structure determination. Nat Methods 14: 290–296

Punjani A, Zhang H & Fleet DJ (2020) Non-uniform refinement: adaptive regularization improves single-particle cryo-EM reconstruction. Nat Methods 17: 1214–1221

Rajendra E, Oestergaard VH, Langevin F, Wang M, Dornan GL, Patel KJ & Passmore LA (2014) The Genetic and Biochemical Basis of FANCD2 Monoubiquitination. Mol Cell 54: 858–869

Rennie ML, Arkinson C, Chaugule V & Walden H (2022) Cryo-EM reveals a mechanism of USP1 inhibition through a cryptic binding site. bioRxiv: 2022.04.06.487267

Rennie ML, Arkinson C, Chaugule VK, Toth R & Walden H (2021) Structural basis of FANCD2 deubiquitination by USP1-UAF1. Nat Struct Mol Biol 28: 356–364

Rennie ML, Lemonidis K, Arkinson C, Chaugule VK, Clarke M, Streetley J, Spagnolo L & Walden H (2020) Differential functions of FANCI and FANCD2 ubiquitination stabilize ID2 complex on DNA. EMBO Rep 21: e50133

Sato K, Toda K, Ishiai M, Takata M & Kurumizaka H (2012) DNA robustly stimulates FANCD2 monoubiquitylation in the complex with FANCI. Nucleic Acids Res 40: 4553–4561

Sims AE, Spiteri E, Sims RJ, Arita AG, Lach FP, Landers T, Wurm M, Freund M, Neveling K, Hanenberg H, et al (2007) FANCI is a second monoubiquitinated member of the Fanconi anemia pathway. Nat Struct Mol Biol 14: 564–567

Smogorzewska A, Matsuoka S, Vinciguerra P, McDonald ER, Hurov KE, Luo J, Ballif BA, Gygi SP, Hofmann K, D’Andrea AD, et al (2007) Identification of the FANCI Protein, a Monoubiquitinated FANCD2 Paralog Required for DNA Repair. Cell 129: 289–301

Sonego M, Pellarin I, Costa A, Vinciguerra GLR, Coan M, Kraut A, D’Andrea S, Dall’Acqua A, Castillo-Tong DC, Califano D, et al (2019) USP1 links platinum resistance to cancer cell dissemination by regulating Snail stability. Sci Adv 5: eaav3235

Terwilliger TC, Sobolev O V., Afonine P V. & Adams PD (2018) Automated map sharpening by maximization of detail and connectivity. urn:issn:2059-7983 74: 545–559

van Twest S, Murphy VJ, Hodson C, Tan W, Swuec P, O’Rourke JJ, Heierhorst J, Crismani W & Deans AJ (2017) Mechanism of Ubiquitination and Deubiquitination in the Fanconi Anemia Pathway. Mol Cell 65: 247–259

Wang R, Wang S, Dhar A, Peralta C & Nikola P (2020) DNA clamp function of the mono - ubiquitinated Fanconi Anemia FANCI - FANCD2 complex. Nature 580: 278–282

Wang S, Wang R, Peralta C, Yaseen A & Pavletich NP (2021) Structure of the FA core ubiquitin ligase closing the ID clamp on DNA. Nat Struct Mol Biol 28: 300–309

Williams SA, Maecker HL, French DM, Liu J, Gregg A, Silverstein LB, Cao TC, Carano RAD & Dixit VM (2011) USP1 deubiquitinates ID proteins to preserve a mesenchymal stem cell program in osteosarcoma. Cell 146: 918–930

Xu X, Li S, Cui X, Han K, Wang J, Hou X, Cui L, He S, Xiao J & Yang Y (2019) Inhibition of Ubiquitin Specific Protease 1 Sensitizes Colorectal Cancer Cells to DNA-Damaging Chemotherapeutics. Front Oncol 9: 1406

